# CRISPR-mediated centromere fission generates neo-chromosomes with distorted meiotic inheritance in Arabidopsis

**DOI:** 10.64898/2026.07.24.740321

**Authors:** Robin Burns, Terezie Mandáková, Chris Morgan, Stephanie Topp, Nicola Gorringe, Tong Li, Katharine Jenike, Piotr Wlodzimierz, Matthew Naish, Martin A. Lysak, Ian R. Henderson

**Affiliations:** Department of Plant Sciences, Downing Street, University of Cambridge, Cambridge, CB2 3EA, United Kingdom; Central European Institute of Technology, Masaryk University, Kamenice 5, 625 00 Brno, Czech Republic; John Innes Centre, Norwich Research Park, Norwich, NR4 7UH, United Kingdom

**Keywords:** Neo-chromosome, centromere, telomere, CRISPR, meiosis, trivalent, distortion, *Arabidopsis*

## Abstract

Centromeres are essential for chromosome segregation and are epigenetically defined by CENH3/CENP-A nucleosomes. Centromere position along chromosomes varies within and between species, ranging from telocentric to metacentric architectures. Yet, how centromere position influences chromosome inheritance and karyotype evolution remains poorly understood. To reposition the centromere, we used CRISPR-Cas9 to break the centromeric satellite array of *Arabidopsis thaliana* chromosome 3. Fission chromosomes rapidly acquired telomeres, converting a metacentric into two stable telocentric neo-chromosomes. Neo-centromere formation involved genetic restructuring and *de novo* satellite higher-order repeat formation, together with epigenetic remodeling of CENH3 and DNA methylation. Crossing the six-chromosome fission line to five-chromosome wild type produced a meiosis-specific trivalent that mis-segregated and generated aneuploidy. Trivalent recombination doubled through two obligate crossovers, which were shifted towards the telomeres. Inheritance was strongly distorted in favor of the wild type centromere, as telocentrics segregated into inviable monosomic gametes. The reciprocal trisomic gametes were associated with centromere-proximal recombination, demonstrating that crossover position governs trivalent segregation. Distortion further increased when CENH3 was over-expressed, implicating centromere strength in the outcomes of trivalent meiosis. Our results reveal rapid centromere remodeling following karyotype change, and how trivalent centromere architecture distorts inheritance, with implications for hybrid incompatibility and synthetic chromosome design.

## Main Text

Eukaryotic karyotypes vary extensively, yet all chromosomes share conserved features required for stable mitotic and meiotic inheritance. Telomeres are necessary to protect chromosome ends, while centromeres mediate segregation during cell division, via deposition of the histone variant CENP-A/CENH3 and kinetochore assembly^1,7^. As centromere identity is epigenetically specified via CENP-A/CENH3, this allows neo-centromeres to form following deletion or inactivation of native centromeres^8–11^. Epigenetic mobility of CENP-A/CENH3 permits centromere repositioning in the absence of structural rearrangements^12,13^. Further, tethering of CENP-A/CENH3, loading factors, and kinetochore proteins is sufficient to create neo-centromeres^14–18^, demonstrating the importance of centromere epigenetic identity. Despite chromatin level flexibility, centromeres commonly associate with repeated DNA sequence architectures, including satellite repeat arrays, and transposon clusters^19,20^, with a high diversity of primary centromeric sequences observed within and between species^21–24^.

Centromere positioning along chromosomes is also dynamic during evolution. Chromosomes show a range of architectures with centromeres located on an axis between the chromosome ends (telocentric), and the chromosome centre (metacentric)^3,12,25–27^. Centromere repositioning is frequently associated with speciation and can contribute to formation of incompatibility barriers^3–5,12,25–27^. For example, Robertsonian chromosome fusions can pair with the ancestral chromosomes during meiosis, forming trivalents and cause gamete aneuploidy and infertility^4–6^. The fertility costs associated with trivalent meiosis, including production of monosomic gametes, will determine whether fusion and fission chromosomes spread, or are purged, from a population. However, an understanding of interactions between recombination, centromere strength, and chromosome segregation during trivalent meiosis is currently lacking.

To investigate how centromere position influences chromosome inheritance, we targeted CRISPR/Cas9 to the satellite arrays of *Arabidopsis thaliana* (hereafter, Arabidopsis) metacentric chromosome three and induced fission. We characterized the resulting telocentric neo-chromosomes using genome assembly, and CENH3 and DNA methylation profiling, which revealed genetic and epigenetic remodeling of the neo-centromeres, following breakage. Using backcrosses between six and five chromosome Arabidopsis we observed meiotic trivalents, associated with doubled recombination rate, distalized crossovers, and distorted Mendelian inheritance in favor of the wild type metacentric centromere. We show that distorted centromere transmission is enhanced by centromere proximal-crossover locations, as well as by CENH3 overexpression. This implies that recombination and centromere strength govern the segregation outcomes of trivalent meiosis. Our integrated view of trivalent recombination and inheritance has relevance to gene flow and selection in Robertsonian hybrids^4–6^, as well as the design and stability of synthetic neo-chromosomes.

## CRISPR-mediated centromere fission generates stable telocentric neo-chromosomes

The centromere of Arabidopsis chromosome-three comprises a 2.2 megabase (Mb) *CEN178* satellite array, which are organized as two sub-arrays in an inverted orientation (1.33 Mb on the p-side, and 0.88 Mb on the q-side), with the p-side array containing four Athila retrotransposon insertions^22,28^ (**Fig. 1a**). Chromosome-specific sequence polymorphisms in *CEN178* repeats provide the opportunity for targeted disruption of a single centromere^22,28^. We identified a guide RNA, *LRCEN3* (5′-AGGCTTACAAGATTGGGTTG-3′) with 154 perfect target sites confined to *CEN3* and no predicted off-target sites genome-wide (**Fig. 1a** and **Extended Data Fig. 1**), which was cloned into the pYAO-Cas9 binary vector for transformation^29^. We transformed Col-0 plants hemizygous for the centromere-three-flanking fluorescent traffic line *CTL3.9*, which allows quantification of meiotic crossover and inheritance within a ∼8.5 Mb centromere-spanning interval^29–32^, to monitor for changes caused by the *LRCEN3* CRISPR-Cas9 construct. As controls, we transformed a no-guide RNA construct, as well as a catalytically dead dCas9-variant with the *LRCEN3* guide RNA.

**Figure 1.**
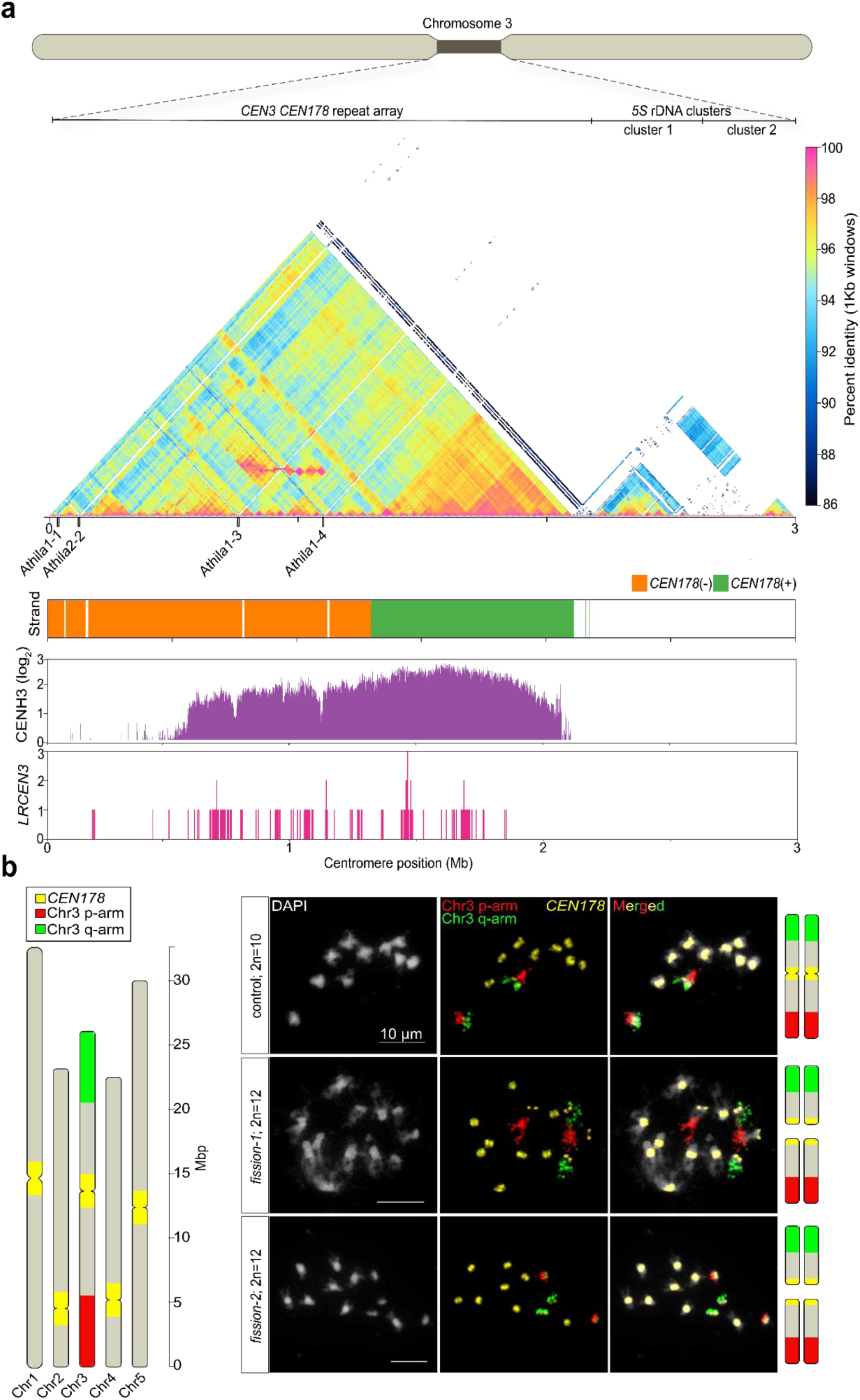
CRISPR-Cas9 targeting centromere 3 satellite repeats in Arabidopsis generates neo-chromosomes. **a.** Sequence identity heat map of chromosome 3 (*CEN3*) using 1 kb windows, with percent sequence identity indicated by the color scale. The ∼2.1 Mb *CEN178* repeat arrays, and two adjacent *5S* rDNA clusters are labelled. Athila retrotransposons (black bars) that are integrated within the *CEN178* repeat arrays are annotated on the x-axis. Below, DNA strand orientation of the *CEN178* repeats are shown, followed by enrichment of CENH3 ChIP-seq enrichment across *CEN3* (log₂[ChIP/input], 1 kb windows). Below, the density of *CRISPR-CEN3* guide RNA target sites are plotted for the guide RNA *LRCEN3*, using 1 kb windows, and permitting no mismatches in the target site. **b.** Schematic representation of the Arabidopsis karyotype, with the arms of chromosome three indicated in red (p arm) and green (q arm), and *CEN178* arrays shown in yellow. Representative micrographs of fluorescently stained mitotic metaphase chromosomes from a control line showing the wild-type Arabidopsis karyotype (2n=10), and from two sibling T_3_ lines carrying chromosome three fission (2n=12) (*fission-1* and *fission-2)*. Chromosomes were counterstained with DAPI (grey), with FISH signals marking the chromosome three p arm (red); q arm (green), and *CEN178* satellite repeats (yellow). In the fission lines, the p-and q-arm signals are spatially separated. Chromosome cartoons adjacent to the photographs depict the inferred chromosome 3 configurations, including separation of the p and q arms into neo-chromosomes in *fission-1* and *fission-2*. Scale bars=10 μm.

Scoring *CTL3.9* meiotic recombination in the *LRCEN3* T_1_ transformants showed both elevated and reduced crossover rates in independent lines, relative to the no-guide controls (Fligner-Killeen *P*=0.027) (**Extended Data Fig. 2** and **Supplementary Table 1**). The *LRCEN3* transformants also exhibited a dwarf phenotype that was absent from no-guide and dCas9 controls (**Extended Data Fig. 3 and 4**). We selected a single *LRCEN3* T_1_ line with significantly elevated *CTL3.9* genetic distance, and ONT sequenced a T_2_ progeny, which revealed reduced read coverage across *CEN3*, but not elsewhere in the genome (**Extended Data Fig. 5**). We performed FISH on 20 self-fertilized T_3_ progeny of this line and identified two siblings with a 2n=12 karyotype in which chromosome-three p– and q-arm signals appeared on separate chromosomes, and which also lacked the *LRCEN3* CRISPR-Cas9 transgene (**Fig. 1b**). These lines were designated *fission-1* and *fission-2* and were advanced through self-fertilization. We observed that the dwarf phenotype co-segregated with the *LRCEN3* CRISPR-Cas9 transgene, but not with the karyotype change, indicating that the developmental changes reflect ongoing CRISPR-Cas9 activity, rather than centromere fission and neo-chromosome presence *per se* (**Extended Data Fig. 3 and 4**).

Oxford Nanopore sequencing and assembly of T_4_ progeny from each fission line yielded contig N50 values >15 Mb and resolved six chromosomes, with chromosome three represented as two distinct molecules (**Fig. 2a** and **Supplementary Table 2**). The two fission lines share nearly identical breakpoints within *CEN3*, consistent with a single ancestral chromosome breakage event (**Fig. 2a**). Read coverage was uniform across the rest of the chromosomes (**Extended Data Fig. 6**). The neo-centromeres are smaller than wild-type *CEN3* (0.89 Mb on neo-3p, and 1.04 Mb on neo-3q) and collectively lack 1,552 *CEN178* repeats relative to the ancestral array, with breakpoints aligning to regions of high *LRCEN3* target site density (**Fig. 2b** and **Supplementary Table 3**). The neo-3p satellite array is largely syntenic with the wild-type p-side array, retaining Athila1-1, Athila1-4, and Athila2-2, but lacking Athila1-3 (**Fig. 2b**). The neo-3q array is partially syntenic with the wild-type q-side but has acquired Athila1-3 and its flanking p-derived *CEN178* repeats, indicating structural translocation between the two sub-arrays during chromosome repair (**Fig. 2b**). Our assemblies are consistent with centromere fission involving coordinated repeat loss and inter-chromosome sequence translocation, with no contribution from the other centromeres (**Extended Data Fig. 7**).

**Figure 2.**
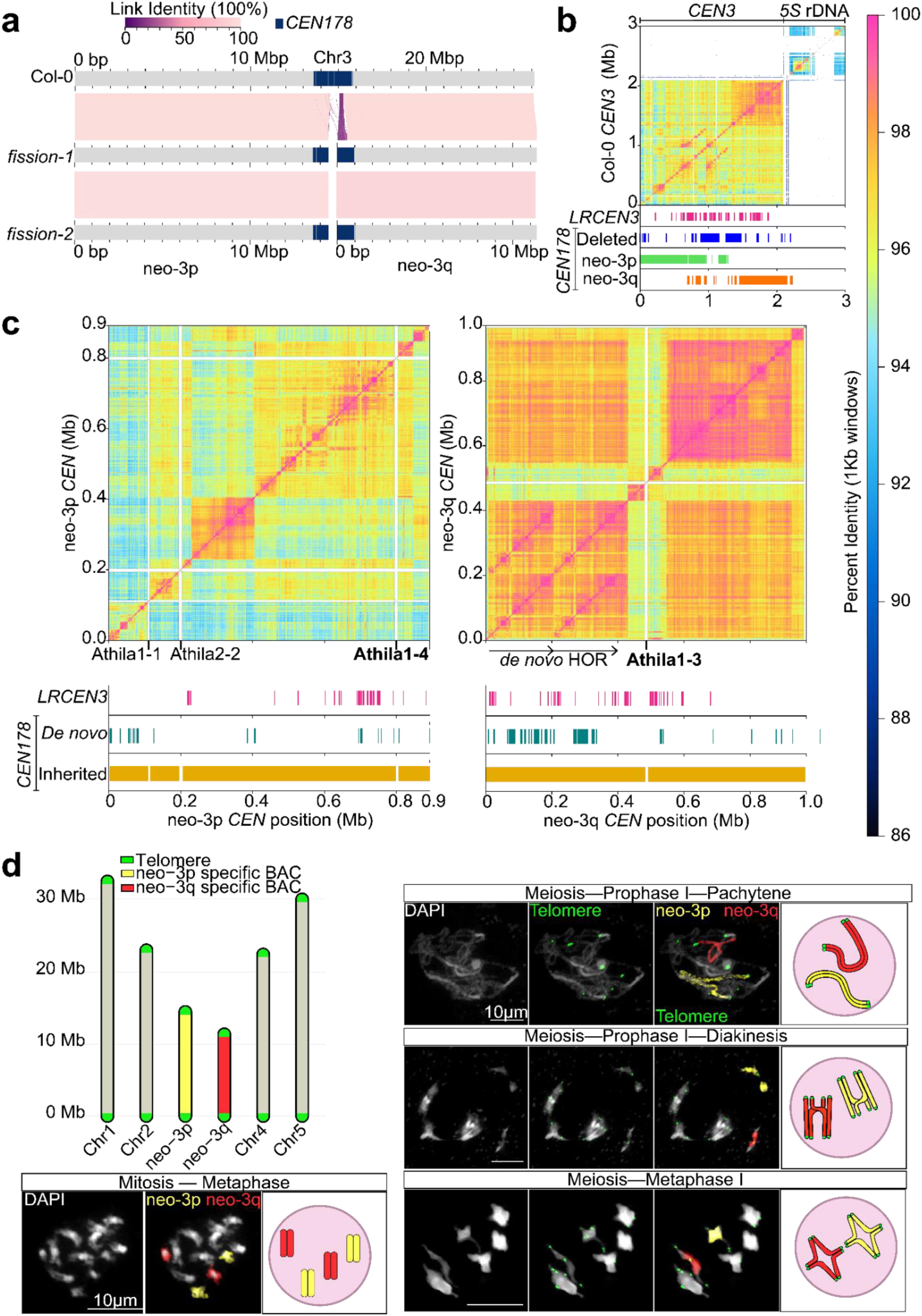
Arabidopsis neo-chromosomes with remodeled centromere satellite arrays and *de novo* telomeres. **a.** Whole-genome alignments comparing Arabidopsis Col-0 wild type chromosome 3 with the two CRISPR-induced fission line assemblies; *fission-1* and *fission-2*. The fission lines share a similar breakpoint within the *CEN3 CEN178* satellite arrays, consistent with shared ancestry. Alignments show that rearrangements are restricted to the centromeric region, whereas the chromosome arms remain syntenic with wild type. The link identity represents percent sequence identity between aligned regions, with light pink indicating higher identity and dark purple indicating lower identity. *CEN178* arrays are indicated by navy blue shading. **b.** Pairwise sequence-identity analysis of the wild type *CEN3* centromeric region. The heatmap represents percent sequence identity calculated in 1 kb windows using ModDotPlot^36^. Tracks below the wild type *CEN3* alignment indicate *LRCEN3* CRISPR target sites (pink), deleted *CEN178* segments from the neo-chromosomes (blue), and *CEN178* sequences corresponding to the neo-centromere arrays, with neo-3p-specific sequences shown in green and neo-3q-specific sequences shown in orange. The positions of Athila retrotransposons are indicated along the x-axis. **c.** Expanded sequence-identity views of the neo-3p and neo-3q centromeric arrays. Heatmaps represent percent sequence identity calculated in 1 kb windows. The remaining *LRCEN3* CRISPR target sites are shown beneath each array in pink. *De novo CEN178* sequences that are not observed in wild type are indicated in teal, and *CEN178* sequences inherited from Col-0 wild type are shown in gold. The *de novo* higher-order repeat on neo-3q is approximately 392 kb and is composed of two approximately 196 kb tandemly duplicated blocks, indicated by arrows. Athila retrotransposons are indicated along the x-axis; neo-3p retains Athila1-1, Athila2-2, and Athila1-4, whereas neo-3q has acquired Athila1-3 via translocation. **d.** Cytological analysis of chromosome 3 fission lines with a haploid chromosome number of n=6. A schematic representation illustrates chromosome 3 split into neo-3p (yellow) and neo-3q (red), with telomeres indicated in green. Representative mitotic metaphase chromosome spreads show DAPI-stained chromosomes in grey, with neo-3p– and neo-3q-specific FISH signals in yellow and red, respectively, located on separate chromosomes; telomeres were not visualized at this stage. Representative meiotic nuclei showing DAPI-stained chromatin in grey, telomeres labelled in green, and neo-3p– and neo-3q-specific signals labelled in yellow and red, respectively, revealing the two fission products as distinct chromosomes that acquired telomeres *de novo*. Adjacent cartoons depict the corresponding chromosome configurations observed. Scale bars=10 μm.

## Neo-centromeres acquired telomeres and a *de novo* satellite higher order repeat

We inspected the neo-centromere arrays for further evidence of DNA sequence restructuring associated with chromosome fission. The neo-3q satellite array contains a ∼392 kb terminal satellite higher-order repeat (HOR) that is absent from the wild type *CEN3*, composed of two ∼196 kb tandemly duplicated blocks (**Fig. 2c**, arrows). The satellite HOR includes 180 *de novo CEN178* repeats (178 of which are unique), compared with 47 new repeats on neo-3p (**Supplementary Table 3**), accounting for most of the new *CEN178* sequence in the fission lines. As both fission lines share this HOR structure, we infer that it formed by at least the T_2_ generation. The neo-3q *de novo* HOR duplication is consistent with homology-driven repair, for example gene conversion or unequal exchange between sister chromatids, following CRISPR-induced breakage. The neo-3q HOR closely resembles structures previously observed in natural *Arabidopsis* pan-centromere variation^22,28^, and in human α-satellite arrays^21^, consistent with DSB-driven repair and recombination seeding the emergence of HORs in satellite centromeres^33,34^.

Stable mitotic and meiotic inheritance of the neo-chromosomes implies that the fissioned ends had been capped by telomeres. Indeed, telomeric repeat arrays were detected at the fission breakpoints of both neo-3p and neo-3q assemblies, spanning ∼2 kb of canonical 5′-TTTAGGG-3′ repeats, that were comparable in length to native telomeres^35^ (**Extended Data Fig. 8**). In contrast to native telomeres, which transition gradually through degenerate variants into sub-telomeric sequence, the neo-telomeres show a sharper boundary between perfect telomeric repeats and internal DNA (**Extended Data Fig. 8**), consistent with recent telomerase action on the broken chromosome ends. The *de novo* telomeres were acquired rapidly within 2–3 generations of the centromere fission event. FISH on meiotic prophase nuclei confirmed telomeric signals at the fission ends of both neo-chromosomes (**Fig. 2d**). We observed that neo-3p and neo-3q segregated independently as bivalents at meiotic metaphase I (**Fig. 2d**), demonstrating the inheritance behavior of true chromosomes.

Short-read sequencing and FISH on independent T_5_ progeny confirmed mitotic and meiotic stability of the 2n=12 karyotype, with no evidence of aneuploidy or copy-number variation (**Extended Data Fig. 9**). RNA-seq of leaves and flower buds detected only single-gene differences between the fission lines and wild-type (**Extended Data Fig. 10**), and self-fertilized fission plants showed normal vegetative growth, pollen viability, and seed set (**Extended Data Fig. 11**). Therefore, targeted DSBs within a native centromere generated two functional neo-chromosomes that acquired telomeres *de novo*, with structural remodeling of centromere satellite arrays and seeding of a novel higher-order repeat. This occurred rapidly within one to two generations between CRISPR transformation and progeny propagation, with minimal transcriptomic or fitness cost when the neo-chromosomes are homozygous.

## Epigenetic remodeling of neo-centromere identity

To test whether the reduced neo-3p and neo-3q arrays support centromeric chromatin, we performed CENH3 ChIP-seq in *fission-2* (**Fig. 3a**). Despite their reduced size, both neo-centromere arrays accumulate CENH3 to levels comparable to the other endogenous centromeres in the same line (**Extended Data Fig. 12**). However, the distribution of CENH3 within the neo-arrays differs from wild-type *CEN3*. In wild-type, CENH3 is enriched on the q-proximal sub-array. In neo-3q, this distribution is preserved, with the neo-3q array derived from sequences that are highly occupied in wild-type, which remain CENH3-occupied in the neo-centromere (**Fig. 3a**). In contrast, in the neo-3p centromere, CENH3 now occupies *CEN178* sequences that show minimal accumulation in wild-type (**Fig. 3a** and **3c**). This demonstrates *de novo* CENH3 deposition on previously unoccupied satellites in the neo-3p centromere, following chromosome fission. The four Athila insertions remain CENH3-depleted relative to the surrounding satellites in the neo-centromeres, as observed in wild-type^22^ (**Fig. 3a**). 3D immuno-FISH confirmed that neo-3p and neo-3q form spatially distinct CENH3 domains on chromosomes, with signal intensities comparable to the endogenous centromeres (**Fig. 3b**). Together this indicates CENH3 redistribution in the neo-centromere satellite arrays, with greatest remodeling in neo-3p.

**Figure 3.**
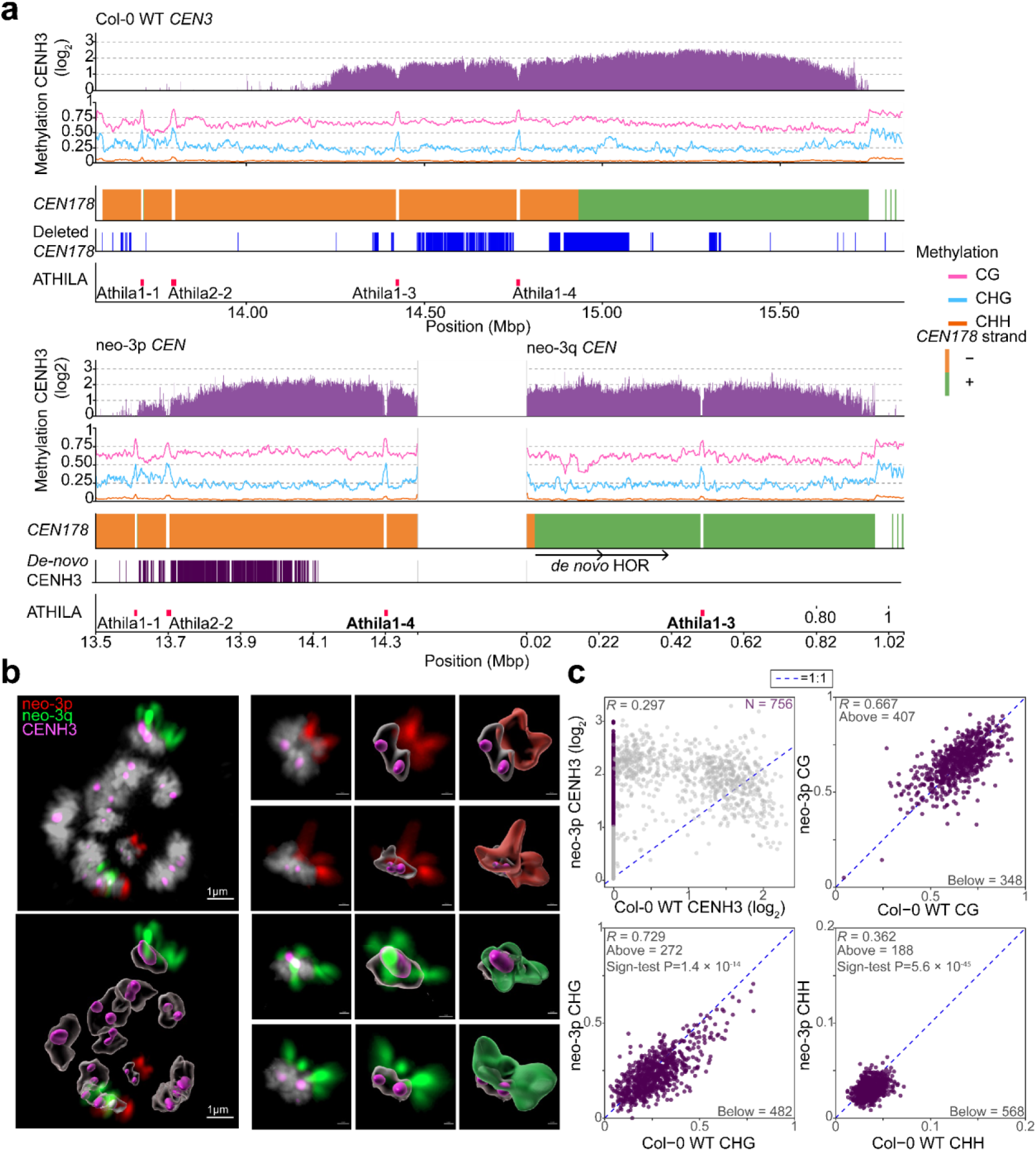
Epigenetic remodeling of neo-centromere chromatin following chromosome fission. **a.** CENH3 ChIP-seq enrichment (purple bars; log₂ChIP/Input) and DNA methylation in 1 kb windows in CG (pink), CHG (blue), and CHH (orange) sequence contexts in wild type Col-0 *CEN3* (top), and the neo-3p and neo-3q centromeres (bottom). Tracks below indicate *CEN178* satellite orientation (+/– strand=green and orange), positions of *CEN178* sequences that are absent from the neo-centromeres (dark blue), *CEN178* with *de novo* CENH3 occupation in neo-3p (dark purple bars), and Athila retrotransposon insertions (red). Athila insertions are epigenetically marked by local elevations in DNA methylation relative to the surrounding *CEN178* satellite sequences and reduced CENH3 occupancy. The physical gap between neo-3p and neo-3q is indicated by the break in the x-axis. **b.** 3D immuno-FISH of mitotic chromosomes showing the spatial organization of fission-derived chromosomes and centromeres. DNA is counterstained with DAPI (grey), CENH3 immunostaining (magenta), and chromosome-arm specific FISH probes label neo-3p (red) and neo-3q (green). Individual chromosomes and their associated CENH3 domains are shown as raw signal projections and as corresponding 3D surface renderings generated from volumetric image segmentation, highlighting spatial relationships between neo-chromosome arms and centromeric chromatin. Scale bars=1 µm. **c.** Scatter plots compare DNA methylation levels between wild-type Col-0 and fission-derived neo-centromeres for CG, CHG, and CHH sequence contexts. Data was generated from the *fission-2* neo-chromosome line. Each point represents a shared *CEN178* repeat between wild type and *fission-2*. Only the DNA methylation of *CEN178* sequences with CENH3 enrichment in *fission-2* (log2[ChIP/Input]≥1), and no detectable enrichment in Col-0 (log2[ChIP/Input]=0), was analyzed (dark purple points). In the scatter plots, the blue dashed lines indicate a 1:1 relationship between wild type and *fission-2* DNA methylation. Pearson correlation coefficients (*R*) are shown for each comparison, together with the number of points above and below the 1:1 diagonal line. One-sided exact sign-test results are shown, *P* values were adjusted using Benjamini–Hochberg correction. CHG and CHH context DNA methylation were significantly reduced in neo-3p relative to Col-0 (*P*=1.4×10⁻¹⁴ and 5.6×10⁻⁴⁵, respectively), whereas CG methylation was not significantly changed (*P*=0.986).

In wild-type *Arabidopsis*, the CENH3-occupied satellite arrays are DNA hypomethylated in non-CG contexts relative to the surrounding pericentromeric heterochromatin^22,28,31^. This is attributed to H3 replacement by CENH3 and consequent depletion of H3K9me2, which is required for maintenance of CHG and CHH DNA methylation^37,38^. Therefore, we predicted that *de novo* CENH3 deposition in the neo-3p satellite array would impose the same epigenetic remodeling of DNA methylation in the newly occupied *CEN178* repeats. Indeed, the *CEN178* satellites that gained CENH3 in neo-3p (log_2_ ChIP/Input ≥1 in *fission-2* versus zero in wild-type) showed depleted CHG and CHH context DNA methylation relative to wild-type, while CG methylation was largely unchanged (one-sided exact sign test, BH-adjusted *P* values: CG=0.986, CHG=1.4×10⁻¹⁴, CHH=5.6×10⁻⁴⁵) (**Fig. 3c**). Therefore, the neo-3p centromere has undergone epigenetic remodeling with both changed CENH3 occupancy, and a decrease in non-CG DNA methylation.

## Meiotic trivalents adopt balanced and unbalanced segregation configurations

To examine the meiotic consequences of combining the ancestral and neo-chromosome fission karyotypes, we crossed the n=6 fission lines to wild-type n=5 plants and analyzed the resulting F_1_ hybrids (n=5×n=6). Self-fertilized fission plants produced the expected n=6 CENH3 foci at pachytene with full homolog synapsis, and neo-3p and neo-3q segregating independently; whereas wild type Col-0 showed five CENH3 pachytene foci (**Fig. 4a** and **Extended Data Fig. 13**). The n=5×n=6 F_1_ hybrids displayed five pachytene CENH3 foci, consistent with formation of a single meiotic trivalent involving the intact wild-type chromosome three and both of the neo-chromosomes (**Fig. 4a**). FISH using chromosome three p– and q-arm probes confirmed trivalent formation at meiotic pachytene, diakinesis, and metaphase I (**Fig. 4b**), while mitotic spreads of five independent F_1_ n=5×n=6 hybrids consistently showed 2n=11 (**Extended Data Fig. 9**), confirming that the trivalent is a meiosis-specific structure.

**Figure 4.**
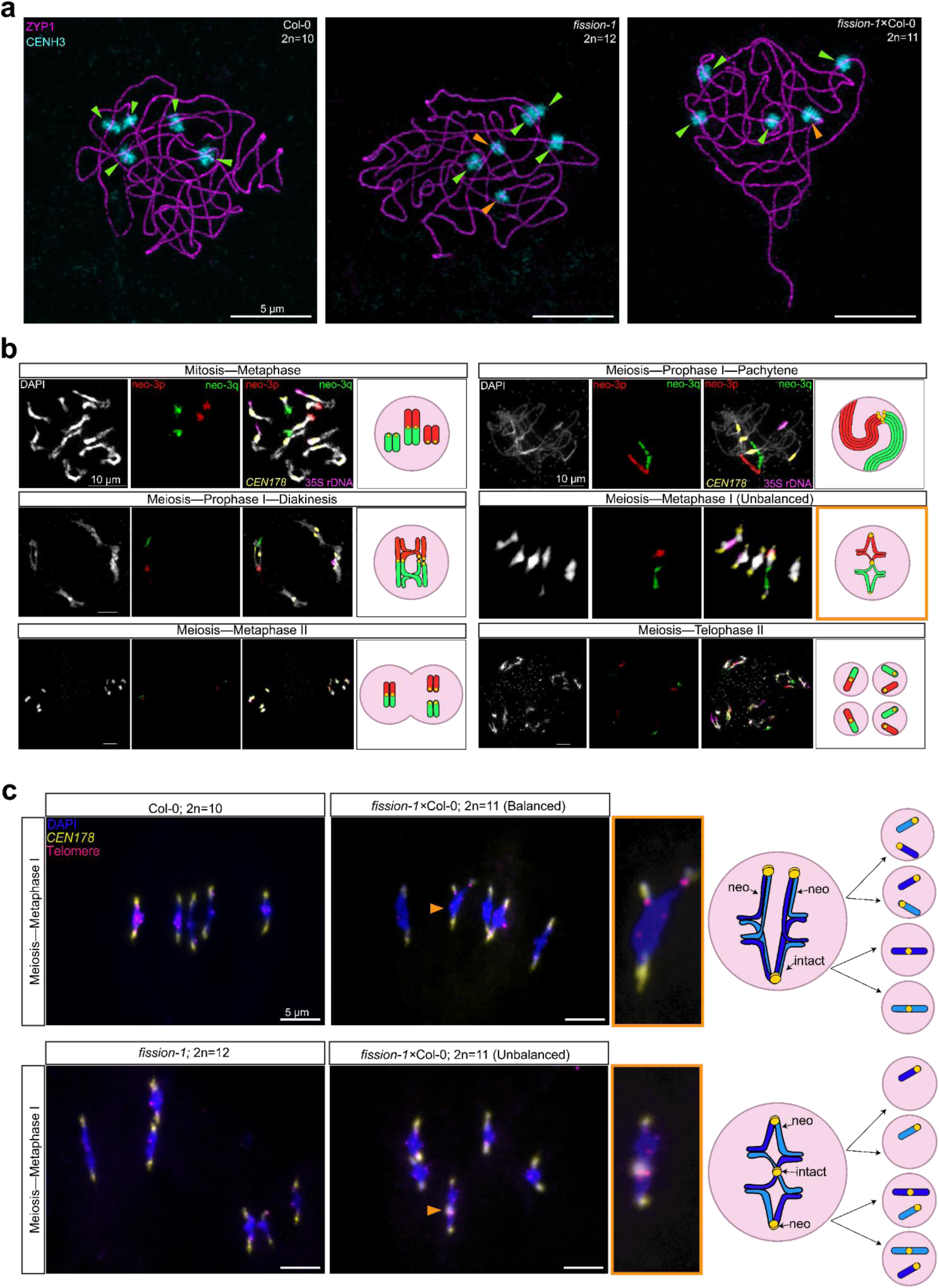
Neo-chromosome meiotic trivalents with balanced and unbalanced segregation at meiosis-I. **a.** Immunostaining of meiotic pachytene chromosomes showing ZYP1 (magenta) and CENH3 (cyan) in Col-0 wild type (left; 2n=10), *fission-1* (middle; 2n=12), and the *fission-1×*Col-0 (n=5×n=6) F_1_ hybrids (right; 2n=11). Arrowheads indicate centromere associations and synaptic configurations, with orange arrowheads highlighting CENH3 signal on the neo-chromosomes. In the n=5×n=6 F_1_ hybrid, five CENH3 foci are observed, consistent with formation of a trivalent involving the two neo-chromosomes and the intact wild type chromosome three. **b.** Neo-chromosome lines (n=6) crossed to wild type (n=5) form trivalent structures during F_1_ hybrid meiosis, but not in mitotic cells. Representative mitotic and meiotic chromosome spreads are shown, labeled by FISH with probes for *CEN178* (yellow), *35S* rDNA (purple), and neo-chromosome specific probes (neo-3p in red, neo-3q in green), illustrating chromosome pairing and structural organization across mitotic and meiotic stages. Each panel represents an independent mitotic or meiotic division. During mitosis (metaphase), neo-chromosomes do not associate with the intact chromosome three. In contrast, during meiosis, the neo-chromosomes and the intact chromosome form a single trivalent. At meiosis metaphase I, non-disjunction configurations of the trivalent are observed, indicative of mis-segregation and highlighted by an orange box. Cartoon diagrams adjacent to each figure depict the corresponding chromosome configurations observed at each stage. **c.** Immunostaining of meiotic metaphase I cells (DAPI=blue; telomeres=magenta; *CEN178*=yellow) reveals variability in trivalent orientation and segregation behavior. Trivalents are indicated by the orange arrowheads. Trivalents can undergo balanced disjunction (alternate), with intact chromosome 3 segregating to one pole, and the fission neo-chromosomes segregating to the opposite pole. Alternatively, non-disjunction of the trivalent leads to unbalanced segregation (adjacent), generating aneuploid meiotic products (e.g. co-segregation of the intact chromosome three with one neo-chromosome), or chromosome loss outcomes in which only one of the two neo-chromosomes is inherited. These chromosome segregation defects are expected to be deleterious due to aneuploidy. Cartoon diagrams of the inferred chromosomes adjacent to the immunostaining micrographs represent the observed trivalent configuration, and the expected euploid and aneuploid gametes. Light and dark blue shading of the chromosomes is used to denote the sister chromatids.

At meiotic metaphase I, F_1_ trivalents adopted two segregation configurations (**Fig. 4b** and **4c**). First, a balanced (alternate) orientation in which the intact chromosome and the two neo-chromosomes segregate to opposite poles, which is predicted to produce euploid gametes (**Fig. 4b** and **4c**). Second, an unbalanced (adjacent) orientation in which the intact chromosome co-segregates with one neo-chromosome, which is predicted to produce monosomic and trisomic outcomes (**Fig. 4b** and **4c**). Unbalanced segregation orientations were not detected in wild-type n=5 or homozygous fission n=6 meiosis (**Fig. 4c**). Consistent with these observations, n=5×n=6 F_1_ hybrid plants showed ∼30% pollen non-viability and ∼20% seed abortion (**Extended Data Fig. 11**). Further, sequencing of 96 F_2_ progeny from *fission-2*×Ler-0 F_1_ detected trisomy of neo-3q (10/96) and neo-3p (3/96), but no double trisomy, or monosomy (**Extended Data Fig. 14**). This is consistent with monosomic gametes or embryos arising from unbalanced trivalent segregation outcomes being inviable. A Col-0×Ler-0 control population showed only one segmental trisomy event in 96 plants (**Extended Data Fig. 15**).

## Trivalents experience doubled meiotic crossovers and distorted fission centromere inheritance

Marey maps of meiotic recombination in the F_2_ population revealed elevated chromosome-three crossover rate in the fission n=5×n=6 hybrid cross, relative to wild type controls (one-sided permutation test *P*=0.001), with no significant change on the other chromosomes (**Fig. 5a** and **Extended Data Fig. 16**). Median per-individual crossover counts on chromosome three doubled from one to two in the n=5×n=6 fission hybrids (Wilcoxon *P*=1.52×10⁻⁴) (**Fig. 5b** and **Extended Data Fig. 17**). Cytological counts of late-pachytene HEI10 foci confirmed that chromosome three recombination doubled in fission n=5×n=6 hybrid meiosis, with trivalents and the summed neo-bivalents both showing significantly more HEI10 foci than wild-type chromosome-three bivalents^39^ (Wilcoxon *P*=5.68×10⁻⁶ and 5.85×10⁻⁴) (**Fig. 5d, 5e** and **Extended Data Fig. 21**). This is consistent with each neo-chromosome in the trivalent receiving an obligate crossover and being regulated as two independent bivalents. Trivalent crossovers were also distally redistributed, and the region of centromere-proximal suppression expanded, relative to the wild-type controls (**Fig. 5a** and **Extended Data Fig. 18**). FTL crosses confirmed remodeling of the recombination landscape, with the distal *420* interval showing elevated crossovers in the trivalent hybrids, while the centromere-spanning *CEN3* and *LTL3.4* intervals were relatively crossover-suppressed, compared to controls (Wilcoxon rank sum test; control vs. *fission-1 P*=2.36×10⁻⁶; *fission-2 P*=1.39×10⁻⁶) (**Extended Data Fig. 20**).

**Figure 5.**
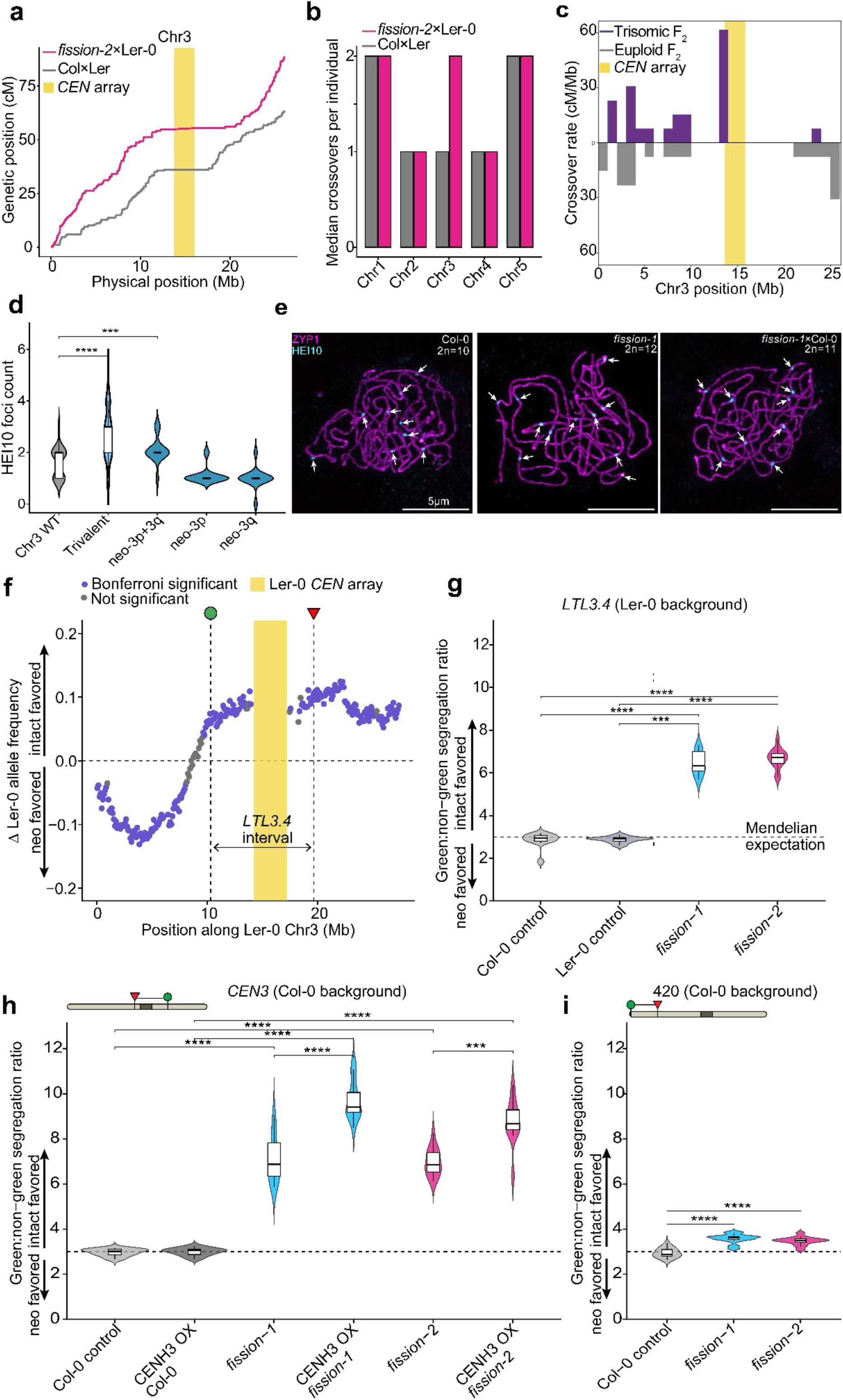
Remodeling of the trivalent meiotic crossover landscape and distorted centromere inheritance. **a.** Marey map of chromosome three showing genetic position (cM) plotted against physical position (Mb) in control (Col-0×Ler-0) and *fission-2*×Ler-0 F_2_ populations. The position of the *CEN178* centromere array is highlighted in yellow. **b.** Median crossover number per individual for chromosomes one to five in control (Col-0×Ler-0) and *fission-2*×Ler-0 F_2_ populations. Crossover number on chromosome three is approximately two-fold higher in the n=5×n=6 trivalent hybrid relative to control, with no significant change on the other chromosomes. **c.** The location of crossovers in the 13 trisomic individuals (purple) compared to an example of a random subset of euploid individuals from the *fission-2*×Ler-0 F_2_ cross. A significant increase of crossovers in centromere-proximal regions is observed in the trisomic individuals (Permutation test *P*<0.001). Violin plots show HEI10 foci counts on chromosome three at late pachytene for Col-0 bivalents, *fission-1*×Col-0 trivalents, summed *fission-1* neo-3p and neo-3q bivalents, and the individual neo-3p and neo-3q bivalent counts from homozygous fission lines (n=6). Internal boxplots show the median and interquartile range, and observation numbers are indicated below each group. HEI10 foci are significantly elevated on the hybrid trivalents, and on summed neo-3p plus neo-3q bivalents, compared to wild type chromosome three bivalents Wilcoxon test *P*<0.001 (***), *P*<0.0001 (****). **e.** Representative late pachytene chromosome spreads immunostained for ZYP1 (magenta) and HEI10 (cyan, with foci indicated by white arrows) in Col-0 (n=5), *fission-1* (n=6), and *fission-1*×Col-0 (n=5×n=6). Scale bars=5 µm. **f.** Differences in the Ler-0 allele frequency (ΔLer-0) in the *fission-2*×Ler-0 F_2_ population, compared to the control Col-0×Ler-0 F_2_ cross, are plotted across Ler-0 chromosome three in 100 kb non-overlapping windows. The yellow bar highlights the position of the Ler-0 chromosome three centromere satellite array. The dashed vertical lines represent the *LTL3.4* fluorescent traffic line (FTL) interval. The dashed horizontal line indicates no change in allele frequency between the populations. Purple points show Bonferroni-significant windows (Fisher’s exact test); grey points are not significant. The location of the green (circle) and red (triangle) *LTL3.4* fluorescent transgenes are indicated above. **g.** Green:non-green fluorescence inheritance ratios in seed from *LTL3.4* heterozygotes with Col-0, Ler-0, *fission-1*, or *fission-2*. The dashed horizontal line marks the expected Mendelian 3:1 inheritance ratio. Blue and pink violin plots represent *fission-1* and *fission-2* hybrid genotypes; grey violins represent controls. Violin plots show the distribution across replicate populations. Internal box plots indicate the median and interquartile range. Wilcoxon test *P*<0.001 (***), *P*<0.0001 (****). **h.** As for F, but analysing green:non-green inheritance of the *CEN3* FTL interval in crosses with Col-0, Col-0 *CENH3-OX*, *fission-1*, *fission-1 CENH3-OX*, *fission-2*, and *fission-2 CENH3-OX*. **i.** As for F, but analysing green:non-green inheritance of the *420* FTL interval in crosses with Col-0, *fission-1*, and *fission-2*.

To determine whether trivalent meiosis biases parental centromere inheritance, we examined Col-0/Ler-0 allele frequencies across chromosome three in the *fission-2*×Ler-0 F_2_ population, relative to the Col-0×Ler-0 control population (**Fig. 5f**). In 100 kb windows along the Ler-0 reference chromosome, Ler-0 allele frequency increased significantly across the chromosome 3 centromere and the q-arm (Fisher’s exact test, Bonferroni-significant; mean Ler-0 frequency 57% versus 48% across the *LTL3.4* FTL interval, odds ratio 1.43, *P*<2.2×10⁻¹⁶) (**Fig. 5f** and **Extended Data Fig. 22**). The p-arm showed a weak Col-0 inheritance bias, consistent with intervening crossovers uncoupling distal regions from centromere-linked distortion. In trivalent hybrid meiosis, the wild-type centromere is therefore preferentially transmitted at the expense of the fission neo-centromeres. This is predicted, as only neo-chromosomes segregate into monosomic gametes during unbalanced trivalent meiosis, providing a transmission advantage to the metacentric wild-type centromere (**Fig. 4c**). To investigate this further, we examined crossover locations in the 13 trisomic individuals obtained from the *fission-2*×Ler-0 F_2_ population. We observed significantly more centromere-proximal crossovers occurring within 1 Mb of the *CEN3* array in the trisomics, compared to the control F_2_ population (one-sided permutation test *P*<0.001) (**Fig. 5c** and **Extended Data Fig. 19).** This is consistent with centromere-proximal crossovers promoting unbalanced segregation of the trivalent at meiosis-I.

FTL crosses, which allow quantification of Mendelian inheritance^32^, confirmed and extended these results. In *LTL3.4* FTL hybrids with the fission lines, the green:non-green seed ratio increased ∼2.3-fold relative to *LTL3.4*×Col-0 and *LTL3.4*×Ler-0 controls (Wilcoxon tests *P*=3.90×10⁻⁵ and 1.07×10⁻⁴) (**Fig. 5g**), with similar magnitudes of change for red:non-red inheritance (**Extended Data Fig. 20**). This confirms that the intact metacentric centromere over-transmits through hybrid trivalent meiosis. We repeated this result in an otherwise Col-0 background lacking Col/Ler polymorphism, using *CEN3* FTL hybrids, which showed the same direction and magnitude of distortion against the fission centromeres (∼2.4-fold, Wilcoxon tests *P*=1.13×10⁻⁵ and 5.10×10⁻⁶) (**Fig. 5h**). In contrast, when we measured transmission of the distal *420* sub-telomeric FTL interval in the hybrids, we observed far weaker ∼1.1-fold distortion (Wilcoxon tests *P*=7.15×10⁻⁶ and 5.76×10⁻⁶) (**Fig. 5i**). This is further consistent with intervening crossovers progressively unlinking distal regions from the distorting inheritance of the fission centromeres during trivalent meiosis.

We sought to test whether changing the dosage of centromeric chromatin would modify the degree of distorted inheritance in hybrid trivalent meiosis, using a *CENH3* overexpression transgene^40^. In fission n=5×n=6 hybrids that also carried *CENH3-OX*, the *CEN3* FTL green:non-green ratio increased to ∼3.1, a further ∼1.3-fold increase over fission hybrids without the *CENH3-OX* transgene (*P*=7.88×10⁻⁶ and 2.08×10⁻⁵) (**Fig. 5h**). *CENH3* overexpression alone in a wild-type background had no effect on FTL inheritance (**Fig. 5h**). This is consistent with an increased frequency of unbalanced segregation outcomes from trivalent meiosis when CENH3 is overexpressed, where neo-chromosomes are lost via monosomic gamete formation (**Fig. 4b-4c**). The F_2_ sequencing and FTL data together establish that meiotic trivalents in fission×wild-type hybrids exhibit distorted inheritance against the newly formed centromeres, which is associated with centromere-proximal crossover locations, and is also sensitive to CENH3 dosage.

## Discussion

We show that targeting CRISPR-Cas9 DNA breakage within a native centromere rapidly generated two functional neo-chromosomes in Arabidopsis, which involved genetic and epigenetic remodeling. Coincident with neo-chromosome formation, the neo-q centromere satellite array acquired a ∼392 kb higher order repeat, composed of two tandem ∼196 kb duplications. This is consistent with homology-directed repair acting on the immediate products of Cas9 cleavage. HORs of comparable structure are widespread in natural pan-centromere variation in *Arabidopsis*^22^, and account for much of the rapid centromere evolution observed in humans and other primate lineages^21,33^. Therefore, similar DSB-driven repair within satellite arrays may seed the punctuated emergence of new HOR structures in natural populations. Further, rapid telomere addition at the fission neo-chromosome ends, comparable to that reported for synthetic maize neo-chromosomes^41^, suggests that telomere healing is a rapid response to broken plant chromosome ends that can stabilize fission products, and thereby allow robust mitotic and meiotic inheritance. This demonstrates that *de novo* telomere and centromere formation are flexible in Arabidopsis, which has implications for engineering synthetic neo-chromosomes that may rapidly adapt to stable forms during propagation.

In the F_2_ progeny of trivalent hybrids, we did not observe transmission of monosomic gametes, although we did observe individuals that inherited the reciprocally produced trisomic gametes. The chromosome three trisomic F_2_ showed a higher incidence of centromere-proximal crossovers, indicating that these events may favour unbalanced segregation outcomes during trivalent meiosis. Consistently, centromere-proximal crossovers have previously been observed to be associated with meiosis-I aneuploidy in humans and *Drosophila*^42,43^. Further, the Mendelian transmission bias we observe across centromere three in n=5×n=6 trivalent hybrids is modified by CENH3 dosage. This suggests that expanded CENH3 chromatin, as well as centromere-proximal crossovers, lead to higher rates of unbalanced trivalent segregation at meiosis-I. This identifies CENH3 dosage and proximal recombination patterns as potential genetic targets that could modify chromosome stability and balanced transmission in Robertsonian hybrid zones^4–6^. Centromere distortion may also explain why fission chromosomes have not yet been observed in the extensive studies of Arabidopsis natural genetic variation^44^.

Simultaneous CRISPR breaks at sub-centromeric and sub-telomeric sites can stably reduce *Arabidopsis* chromosome number from n=5 to n=4, by translocating chromosome arms and eliminating a centromere^45^. Here, we show that centromere fission generates two new centromeres from one, which is also tolerated when homozygous. However, the fission centromeres are selected against in hybrid meiosis, due to monosomic outcomes of trivalent meiosis involving the neo-chromosomes; whereas the intact chromosome can transmit as a trisomic in the reciprocal gametes. As monosomic gametes abort directly, or cause embryo lethality, this causes distorted inheritance against the fission neo-centromeres. We also show that intervening meiotic crossover events can unlink the distal regions from the distorted neo-centromere inheritance. Therefore, while trivalent meiosis can bias transmission against the neo-centromeres, crossovers in the chromosome arms unlink distal regions from the centromeres, with the potential to permit gene flow in Robertsonian hybrids. Extension of centromere DSB targeting to polysomic polyploids, holocentrics, and lineages with extreme chromosome numbers, may reveal further meiotic mechanisms that shape karyotype evolution and reproductive isolation, as well as inform on synthetic neo-chromosome design.

## Data availability

All sequencing data generated in this study have been deposited in the European Nucleotide Archive (ENA) under PRJEB108614, including: R9 ONT FASTQ files for *LRCEN3* T2-1 and the CRISPR no-guide control; R10 raw unaligned bam with methylation calls for fission-1 and fission-2; R10 raw ONT unaligned bam file with methylation calls included for Col-0 wild-type; RNA-seq FASTQ files from leaf and flower bud tissues of the fission lines and the Col-0 wild type; ChIP-seq FASTQ files for the fission lines; DNA sequencing FASTQ files from the T_5_ generation of the fission lines; and DNA sequencing FASTQ files from the F2 populations used for genetic map construction. Genome assemblies of the fission lines generated from the R10 ONT data have been deposited under GCA_981082055 (*fission-1*) and GCA_981082065 (*fission-2*).

## Author contributions

I.R.H, R.B, and T.M conceptualized and designed the study. I.R.H and R.B. co-supervised the project. Genomic DNA and RNA extractions and genome assembly were performed by R.B. Molecular cloning was performed by R.B and P.W. Plant material was generated by S.T, R.B, and N.G. FISH and immunostaining experiments were performed by T.M and C.M. CENH3 ChIP sequencing and DNA methylation profiling were performed by M.N. R.B and T.L. R.B, T.M, C.M, M.N, N.G, T.L, K.J, M.A.L and I.R.H analysed the data. R.B and I.R.H wrote the paper, with input from all authors.

## Acknowledgements

This work was supported by grants from the European Research Council (AdV-EvoPanCen), and BBSRC-UKRI grant BB/Y009487/1 to IRH, and from the Czech Science Foundation Project No. 24-11371S to TM. RB was supported by a long-term EMBO postdoctoral fellowship (ALTF224-2022) and a Broodbank Fellowship from the University of Cambridge. TL was supported by a CSC fellowship. MN was supported by a Broodbank Fellowship from the University of Cambridge. CM was supported by BBSRC-UKRI and a Royal Society University Research Fellowship. MAL and TM were supported by the project TowArds Next GENeration Crops (TANGENC; CZ.02.01.01/00/22_008/0004581) of the ERDF Programme Johannes Amos Comenius. We thank Andreas Houben and Chris Franklin for supplying primary antibodies used in this study.

## Competing interests

There are no competing interests to declare.

---

**Extended Data Figure 1.**
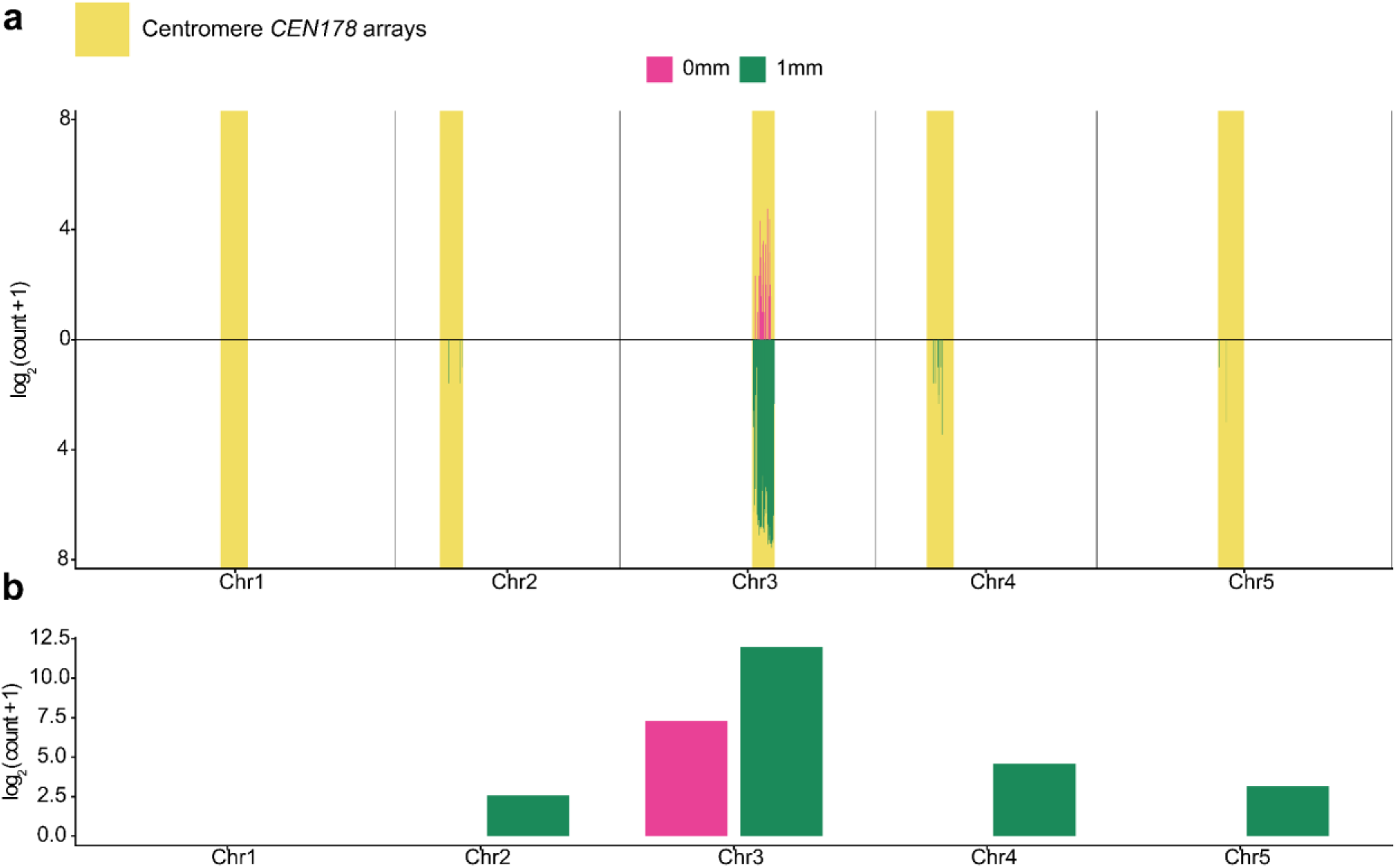
Genome-wide distribution of *LRCEN3* CRISPR target sites. **a.** Binned distribution of *LRCEN3* target sites across concatenated chromosomes (Col-0). Perfect matches (0 mm, pink) are plotted above and single mismatches (1 mm, green) below the axis. Values are shown as log₂(count+1). Centromeric satellite array positions are shaded in yellow. **b.** Per-chromosome counts of *LRCEN3* target sites (0 and 1 mm), plotted as log₂(count+1).

**Extended Data Figure 2.**
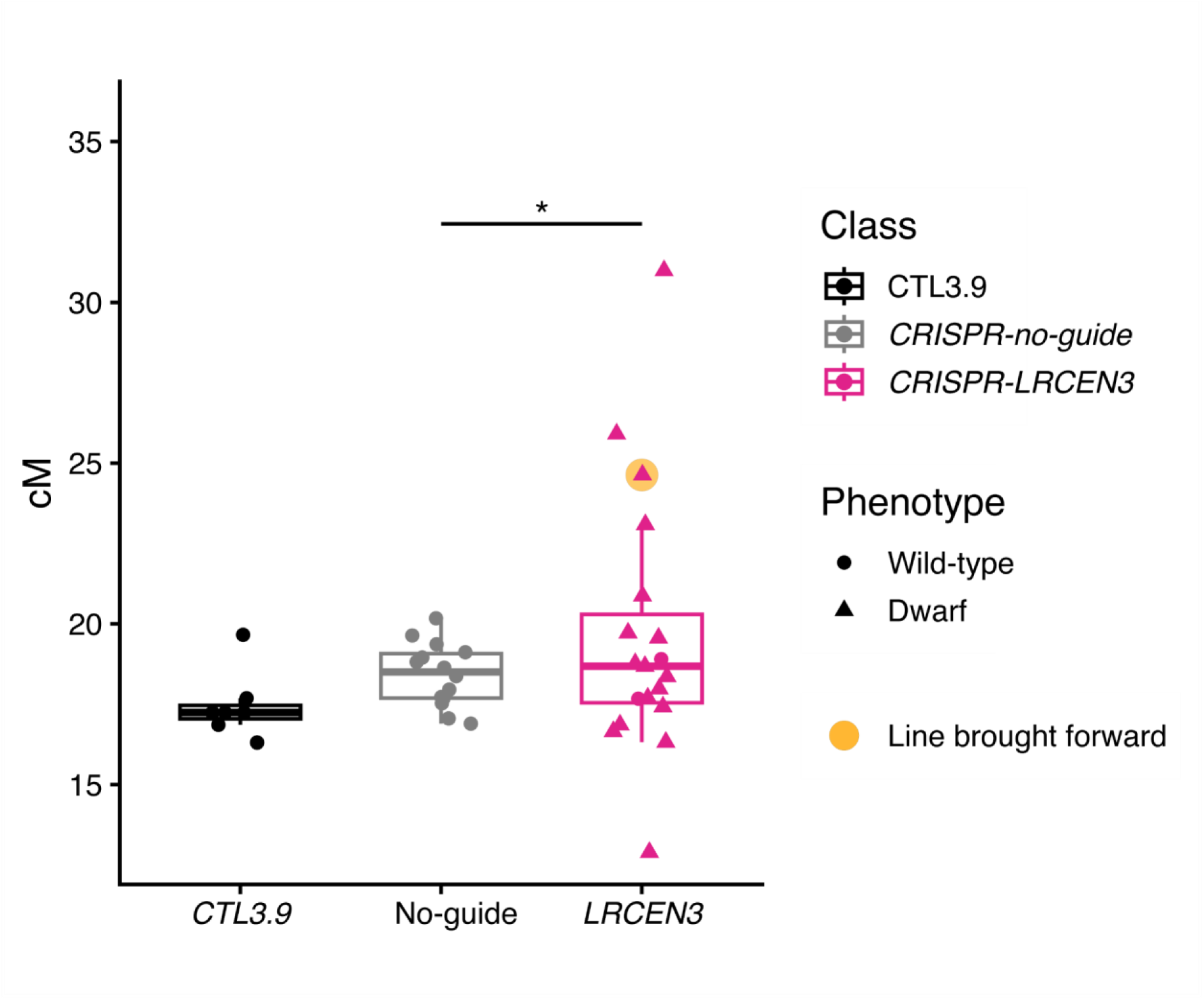
*CTL3.9* crossover frequency in plants transformed with *CRISPR-CEN3* constructs. Boxplots show meiotic crossover recombination rate (centiMorgans, cM) for T_1_ plants across three genotype classes (left to right): untransformed *CTL3.9* (black), no-guide Cas9 (grey), and *LRCEN3* (pink) transformed T_1_ plants. Points represent data from individual plants. Plants with a wild type developmental phenotype are indicated by circles; whereas plants with a dwarf phenotype are indicated by triangles. The orange circle highlights the *CRISPR-LRCEN3* T_1_ line that was continued to T_2_. Increased variance in cM per guide RNA relative to NC was assessed using Fligner-Killeen tests. Only significant comparisons are annotated with bars and asterisks (*). *LRCEN3* vs no-guide, *P*=0.027. The dwarf phenotype incidence by construct were: *CTL3.9*, 0 of 7; no-guide, 0 of 14; and *LRCEN3*, 17 of 19. Boxes show the median and inter-quartile range (IQR), and whiskers indicate 1.5× the IQR.

**Extended Data Figure 3.**
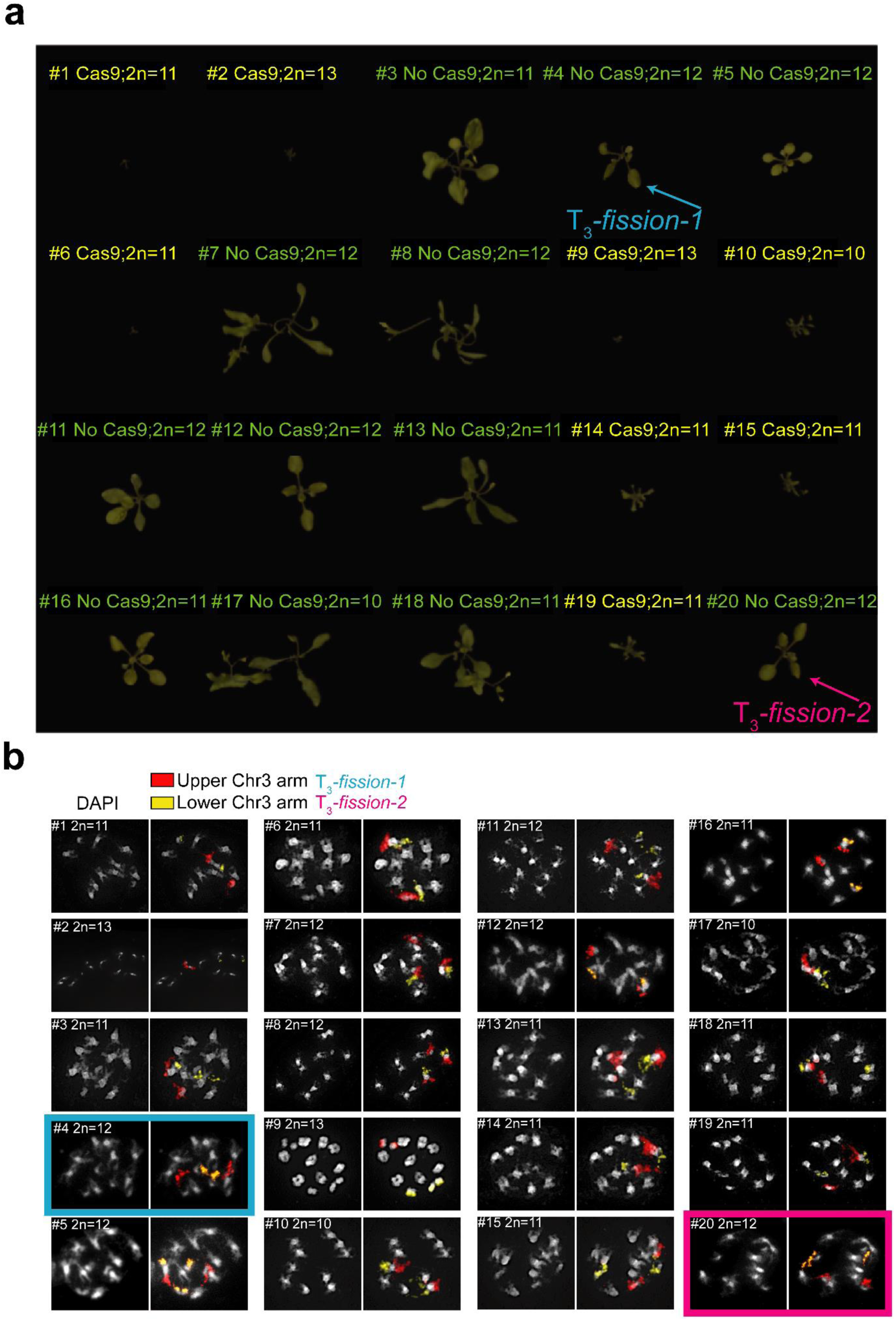
Developmental phenotypes associated with *CRISPR-LRCEN3* transformation. **a.** Twenty representative T_3_ plants derived from a single T_2_ *Cas9-CRISPR-LRCEN3* parent line are shown. The T_3_ individuals segregate for chromosome number (2n=12, 11, or 10) and for presence of the *Cas9-CRISPR-LRCEN3* T-DNA. Cas9-plants lacking the T-DNA are labeled in green. Cas9+ plants retaining the *Cas9-CRISPR-LRCEN3* T-DNA are labeled in yellow. The observed karyotype distribution was: 2n=10 (2 Cas9+, 1 Cas9-), 2n=11 (3 Cas9+, 5 Cas9-), and 2n=12 (3 Cas9+, 6 Cas9-). Plants were imaged 21 days post-germination. Two lines were selected for further analysis, T_3_-*fission-1* (pink) and T_3_-*fission-2* (blue) and are indicated by arrows. **b.** Representative mitotic chromosome spreads used for karyotype determination. DNA was counterstained with DAPI (grey). Fluorescence *in situ* hybridization was performed to label the p arm of chromosome three in red and the q arm in yellow. T_3_-*fission-1* (pink) and T_3_-*fission-2* (blue) are indicated by shaded boxes.

**Extended Data Figure 4.**
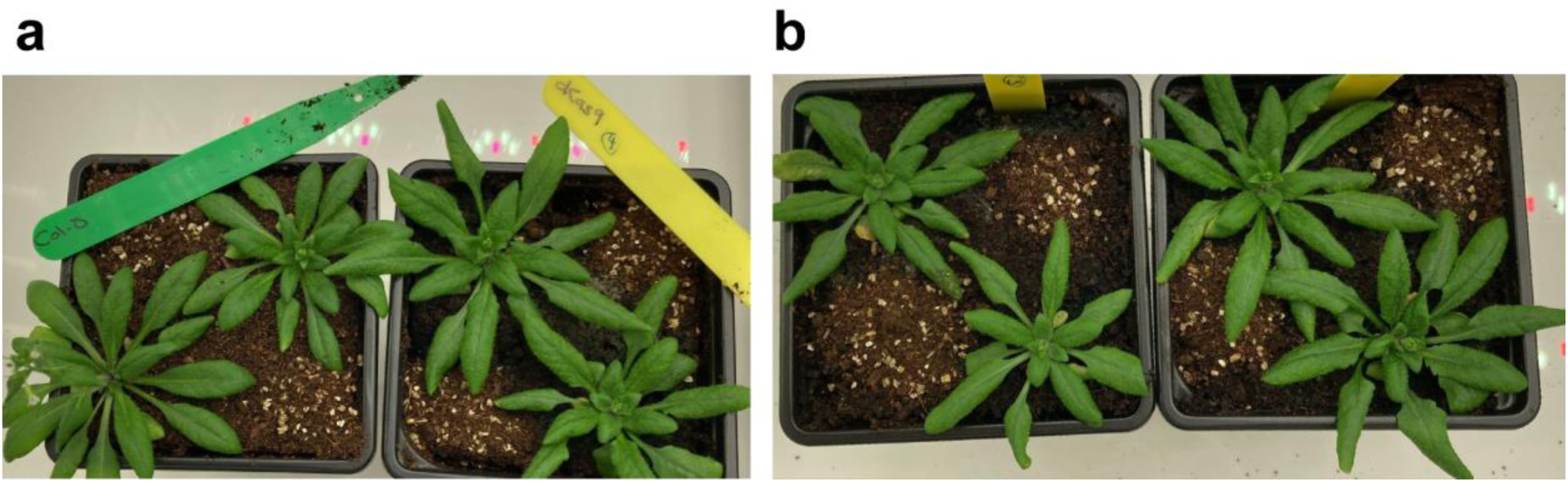
Nuclease-dead *dCas9-CRISPR-LRCEN3* transgenics do not exhibit a dwarf phenotype. **a.** Photograph of wild type (Col-0) plants (left), and *dCas9-CRISPR-LRCEN3* transformants (right). **b.** Photograph of four additional *dCas9-CRISPR-LRCEN3* transgenic plants. Across all plants examined, the vegetative morphology of *dCas9-CRISPR-LRCEN3* individuals was indistinguishable from wild type.

**Extended Data Figure 5.**
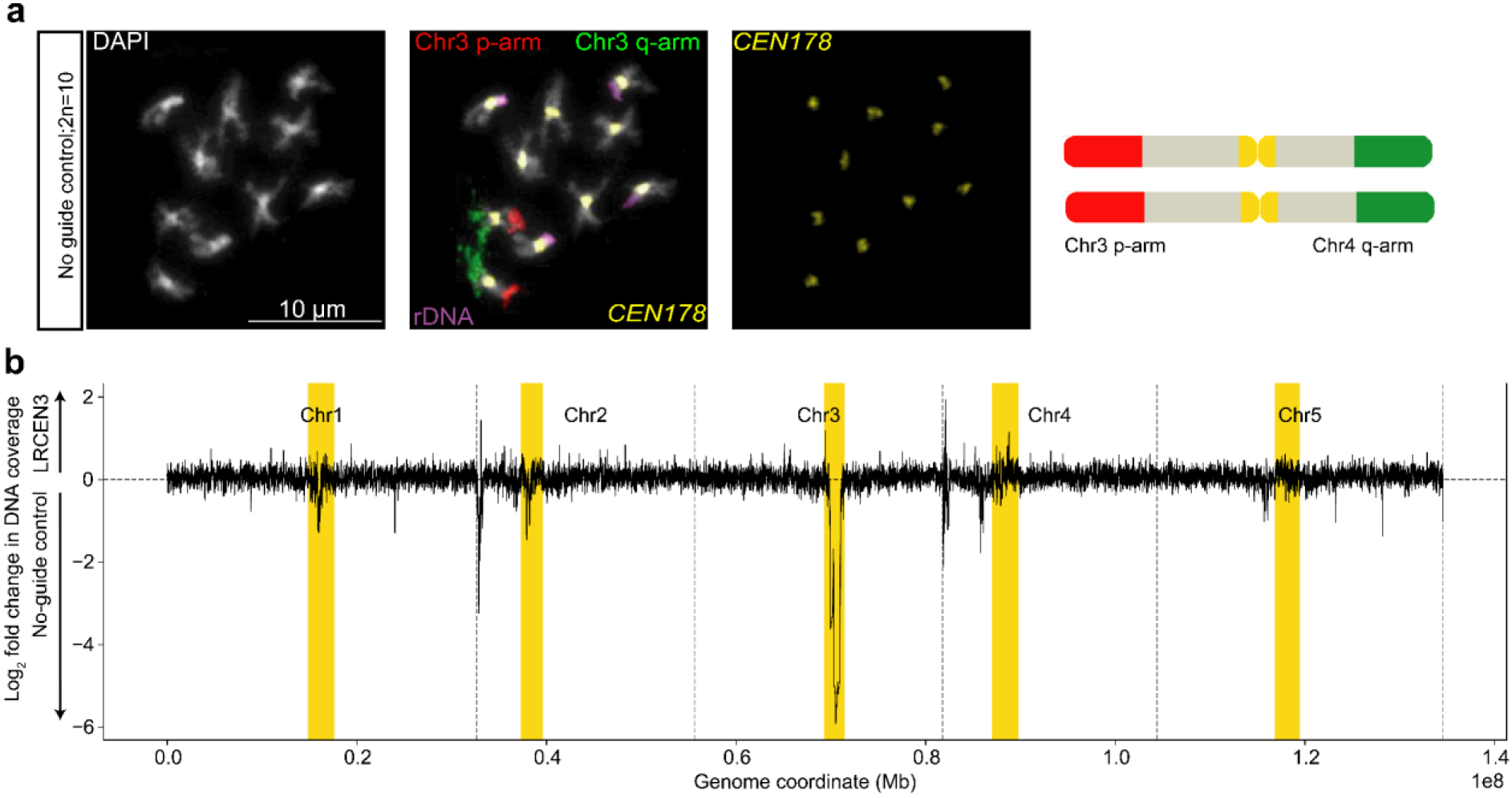
Whole-genome sequencing coverage in the T₂ *Cas9-CRISPR-LRCEN3* line relative to no-guide-Cas9 controls. **a.** Micrograph of FISH staining of the no-guide control showing the wild type Arabidopsis karyotype (2n=10). DAPI staining (white) is shown, as well as probes hybridizing to the chromosome three p arm in red, and q arm in green, 35S rDNA in purple, and *CEN178* in yellow. The cartoon next to the micrograph depicts the intact Chr3 karyotype observed in the images. **b.** Log₂-transformed read depth (*LRCEN3*/no-guide control) is plotted across chromosomes one to five, with vertical dashed lines indicating chromosome boundaries and yellow shading marking centromeric *CEN178* arrays. Values near zero indicate similar coverage between samples, whereas negative values reflect reduced coverage in the *Cas9-CRISPR-LRCEN3* line relative to the control. Sequencing coverage depletion was observed in the centromere of chromosome three, consistent with loss of DNA sequence at this locus, while other centromeric regions were unchanged.

**Extended Data Figure 6.**
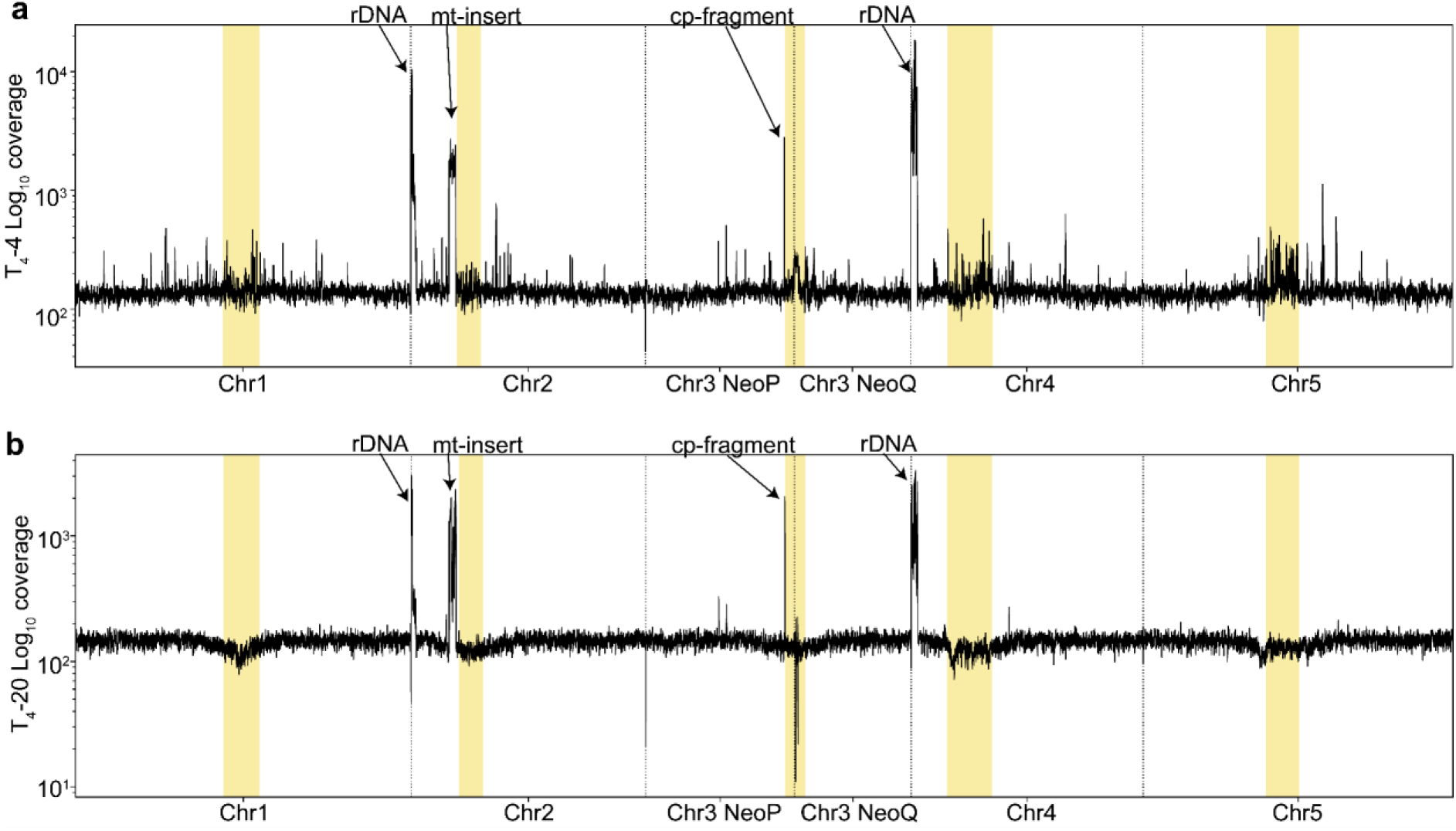
Aligned ONT sequencing read coverage across neo-chromosome fission line genome assemblies. **a.** Black traces show normalized ONT read depth (log_10_ coverage) of *fission-1* across each chromosome aligned to the corresponding fission assembly. Yellow shading indicates centromeric *CEN178* repeat arrays. The fission chromosome assembly exhibits pronounced coverage peaks at ribosomal DNA (rDNA) loci near the beginning of chromosomes two and four, as well as at pericentromeric organellar sequence insertions, including a mitochondrial fragment (mt-insert) on chromosome two, and a chloroplast-derived region (cp-fragment) on chromosome three, indicated by arrows. **b.** As for A., but analysing line *fission-2*.

**Extended Data Figure 7.**
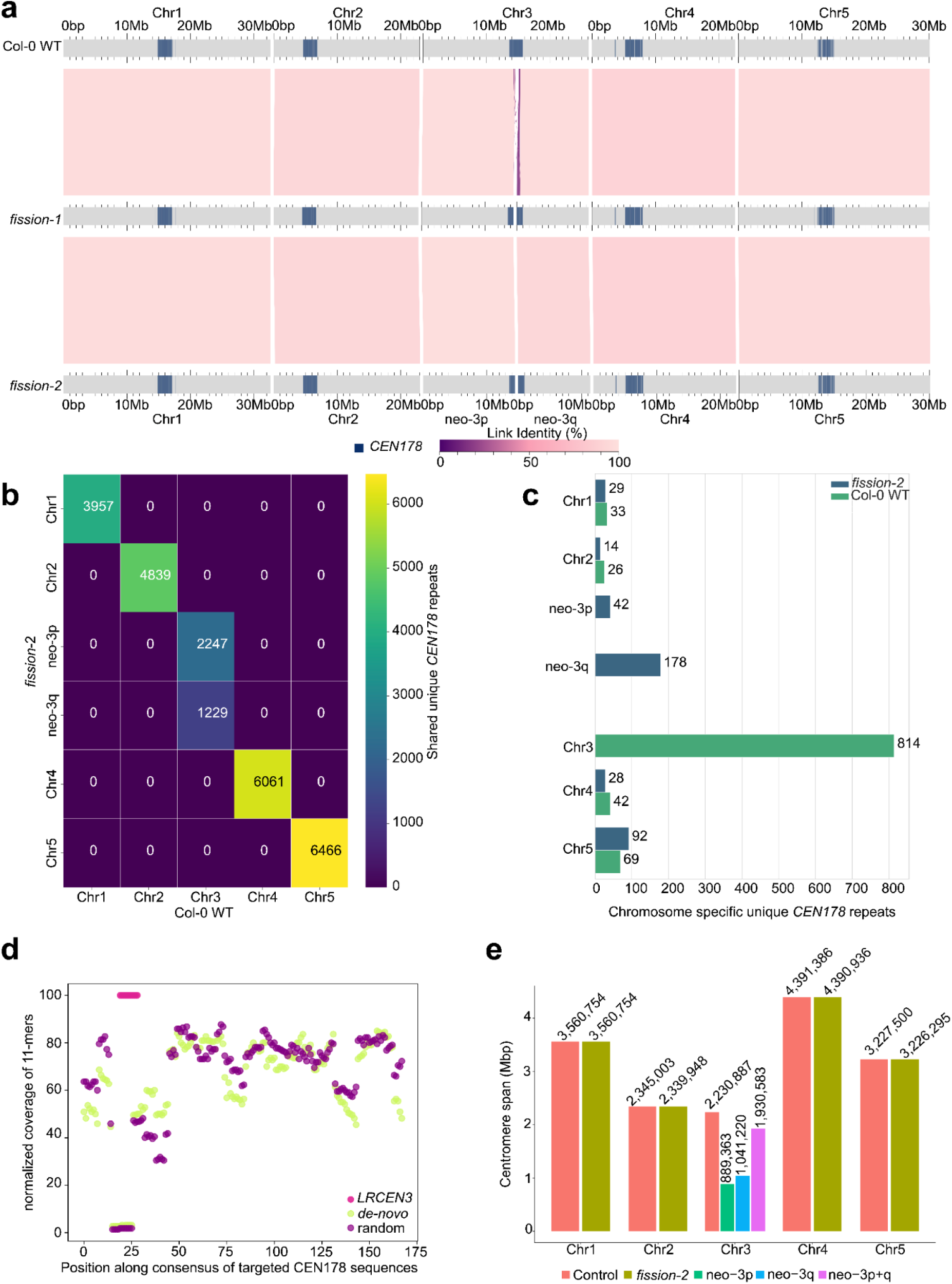
Whole-genome alignment of Col-0 wild type and chromosome three fission genome assemblies. **a.** Whole-genome alignments show that both chromosome three fission lines maintain overall chromosomal synteny relative to Col-0 wild-type. Both *fission-1* and *fission-2* assemblies share a major rearrangement in centromere three, with a similar breakpoint position between the fission lines. Together, these results are consistent with the neo-chromosomes neo-3p and neo-3q in *fission-1* and *fission-2* likely originating from the same centromeric fission event. Sequence assembly alignments were visualized using AliTV (*1*). Centromere *CEN178* arrays are indicated in dark blue shading. **b.** Heatmap showing the number of shared unique *CEN178* repeats between Col-0 wild type chromosomes (x-axis), and those of the *fission-2* neo-chromosome individual (y-axis). We focus on unique *CEN178* sequences as we can assign them to a specific chromosome. The analysis demonstrates that *CEN178* repeats located in the neo-3p and neo-3q centromeres are derived from *CEN3* of Col-0 wild type. **c.** Bar plot showing chromosome-specific unique *CEN178* repeats in Col-0 wild type and *fission-2*. The majority of the chromosome-specific *CEN178* repeats are located in centromere 3 of Col-0 wild type, indicating that a large fraction of *CEN3*-associated repeats were lost following centromere fission and rearrangement. **d.** Normalized 11-mer coverage across a consensus sequence generated from *CEN178* repeats targeted by the *LRCEN3* guide. 11-mers from *de novo* neo-centromere *CEN178* repeats (green) and randomly selected *CEN3* wild-type *CEN178* repeats (purple) are plotted, with the *LRCEN3* target region highlighted in pink. 220 monomers were sampled for each. Both populations show equivalent coverage profiles, including a lack of coverage at the *LRCEN3* target site, meaning this region is not present in both sets. **e.** Centromere array sizes of Col-0 wild-type chromosomes (red) compared to *fission-2* (olive), with neo-3p (green) and neo-3q (blue), and the sum of neo-3p and neo-3q (purple) highlighted. In total ∼300 kb of the original *CEN3* array has been deleted in the fission lines.

**Extended Data Figure 8.**
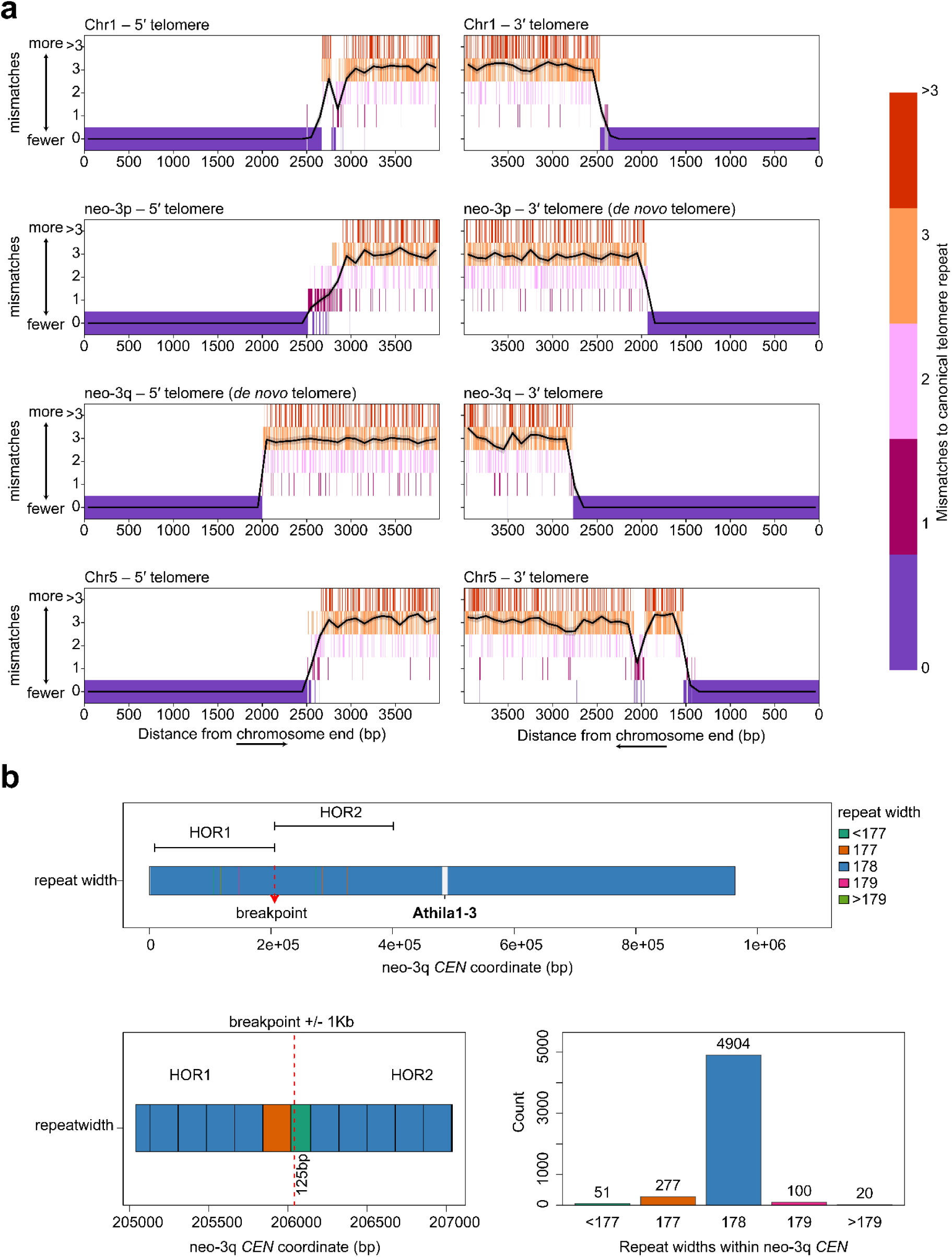
Telomere repeats of neo-chromosomes and break site between HOR1 and HOR2 on neo-3q. **a.** Telomeric sequence composition along the distal 4 kb of chromosome ends of the *fission-2* genome assembly, including chromosomes one, five, and the fission-derived neo-chromosomes neo-3p and neo-3q. For each sliding 7-bp window, mismatch counts compared to the canonical telomeric repeat (5′-TTTAGGG-3′) are plotted. Telomeric termini exhibit a gradual rise in degenerate telomeric variants (1–3 mismatches), before transitioning into non-telomeric sequences (>3 mismatches). In contrast, both neo-3p and neo-3q telomeres display a sharp boundary between perfect telomeric sequence and internal DNA, indicating rapid *de novo* telomere additions that are comparable in size to wild type chromosomes. The black line indicates the mean level of mismatches with the shaded region indicating the 95% confidence interval. Chromosomes two and four telomeres are excluded from the analysis, as the unassembled repetitive ribosomal DNA arrays at their nucleolar organiser regions prevent assembly of the p-arm telomeres. **b.** Structure of the neo-3q centromeric higher-order repeat region surrounding the breakpoint between HOR1 and HOR2. HOR1 spans neo-3q:8526–206040bp, and HOR2 spans Chr3_NeoQ:206041–401185bp. Repeat-width classes are shown across the neo-3q centromeric region, with the breakpoint indicated by a red dashed line. A magnified ±1 kb view of the breakpoint shows the local transition between HOR1 and HOR2 repeat organization, including a 125-bp interval at the junction. The bar plot summarizes repeat-width counts within the neo-3q centromeric region, showing that 178-bp repeats are the dominant repeat class.

**Extended Data Figure 9.**
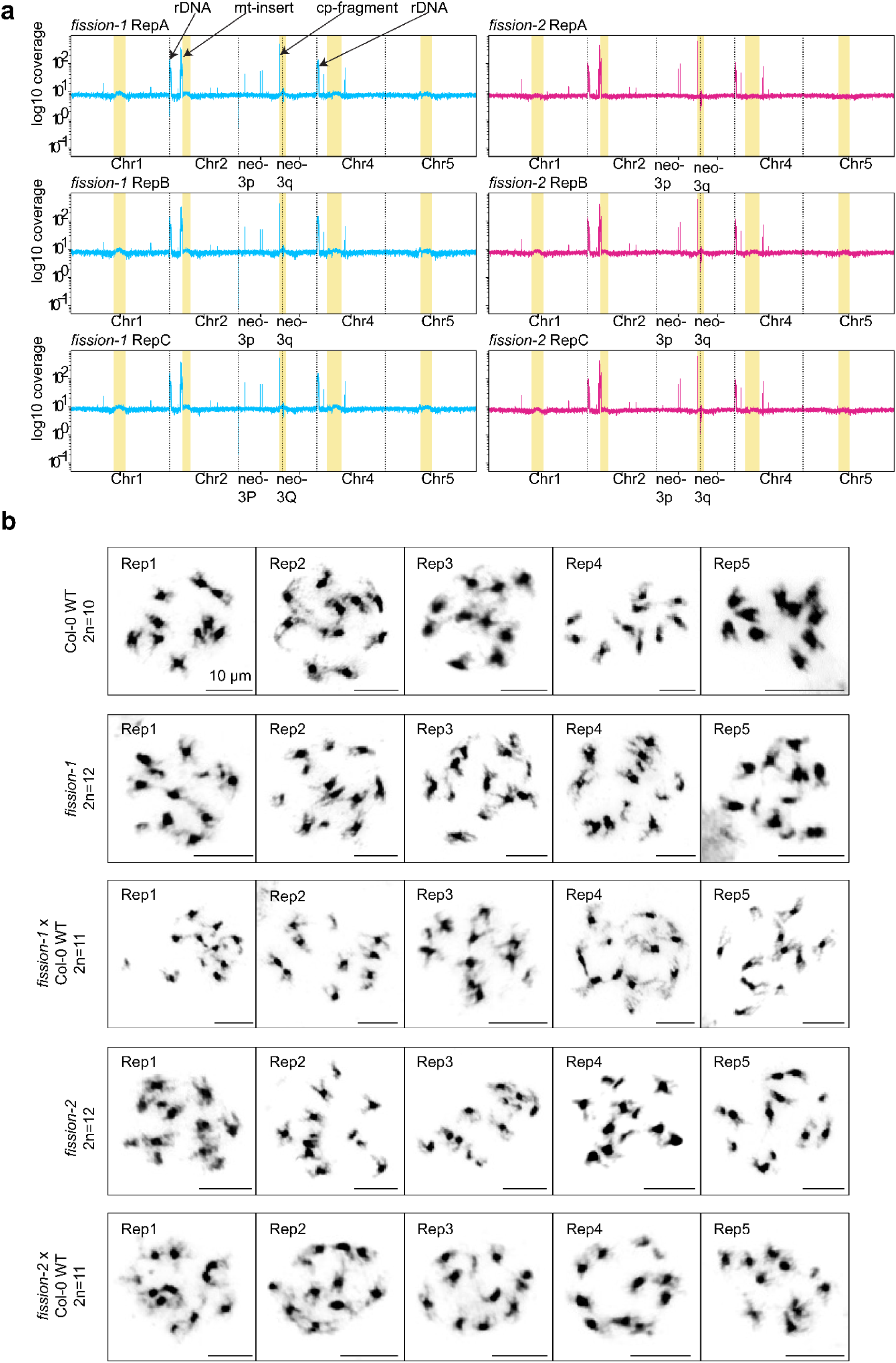
Mitotic and meiotic genome stability of neo-chromosome lines. **a.** Whole-genome short-read sequencing profiles (∼10–15× coverage) of three independent T_5_ progeny from *fission-1* (blue) and *fission-2* (pink) neo-chromosome lines. Illumina read coverage (log_10_ scale) is plotted across all five chromosomes. Yellow shading marks centromeric *CEN178* arrays, and dashed lines indicate chromosome boundaries and neo-chromosome breakpoints (neo-3p and neo-3q). Local high coverage peaks indicated by arrows correspond to known high-copy loci, including the *45S* rDNA arrays (rDNA), as well as a mitochondrial DNA insertion proximal to the chromosome two centromere (mt-insert), and a chloroplast DNA insertion near the neo-3p centromere (cp-fragment). Overall coverage uniformity indicates stable genome structure without large-scale copy-number changes, with no evidence of aneuploidy of neo-3p and neo-3q. **b.** Representative metaphase mitotic chromosome spreads from wild type (Col-0, 2n=10), *fission-1* (2n=12), *fission-1*×Col-0 F_1_ hybrids (2n=11), *fission-2* (2n=12), and *fission-2*×Col-0 F_1_ hybrids (2n=11), with five biological replicate spreads shown for each genotype. Chromosome numbers are consistent with the expected karyotypes, supporting mitotic stability of the neo-chromosomes in somatic cells. Chromosomes were counterstained with DAPI. Scale bars=10 μm.

**Extended Data Figure 10.**
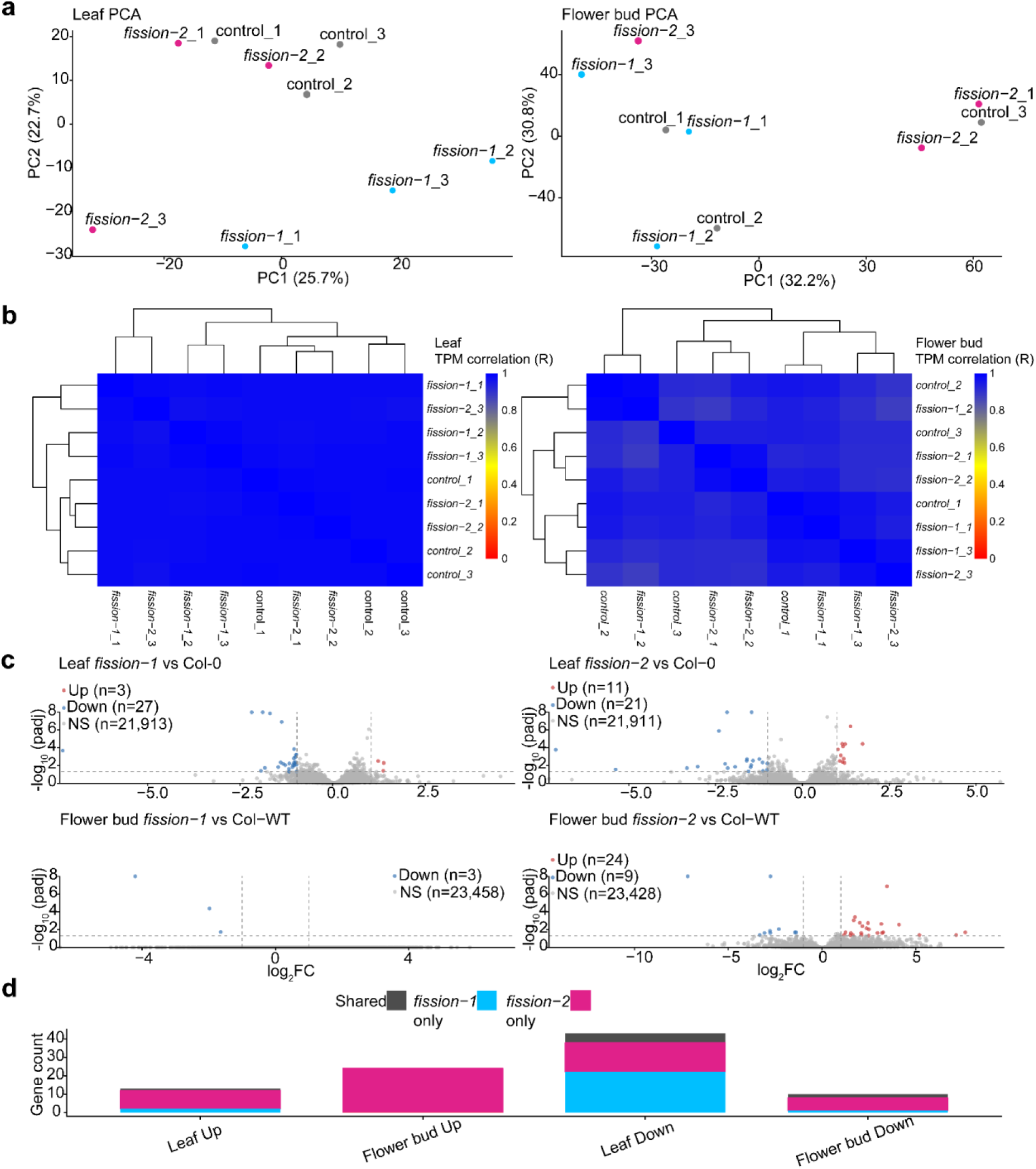
Transcriptomic comparisons between wild type and centromere fission lines. **a.** Principal component analysis (PCA) of log₂(TPM+1) gene expression values from leaf (left) and flower bud (right) tissues. Each data point represents an individual biological replicate from wild type (Col-0), *fission-1*, or *fission-2*. The percent variance explained by each principal component is indicated on the axes. **b.** Hierarchically clustered correlation heatmaps of transcript abundance (TPM) across samples for leaf (left), and flower buds (right). Dendrograms were generated by hierarchical clustering (average linkage) on the sample distance matrix. Colors indicate pairwise Pearson correlation coefficients**. c.** RNA-seq differential expression analysis of *fission-1* and *fission-2* relative to wild type in leaf and flower bud tissues. Volcano plots show log₂ fold change versus −log₁₀(adjusted *p* value) from DESeq2 negative binomial generalized linear models using Wald tests, with Benjamini–Hochberg multiple-testing correction (padj). Dashed lines indicate thresholds of padj<0.05 and |log₂FC| ≥ 1; numbers of up-regulated, down-regulated, and non-significant genes are shown in the legends. For visualization, −log₁₀(padj) values greater than 8 were truncated to 8 on the y-axis**. d.** Overlap of differentially expressed genes (padj<0.05; |log₂FC| ≥ 1) between *fission-1* and *fission-2* within each tissue. Stacked bars show numbers of up– and down-regulated genes partitioned into shared and genotype-specific sets.

**Extended Data Figure 11.**
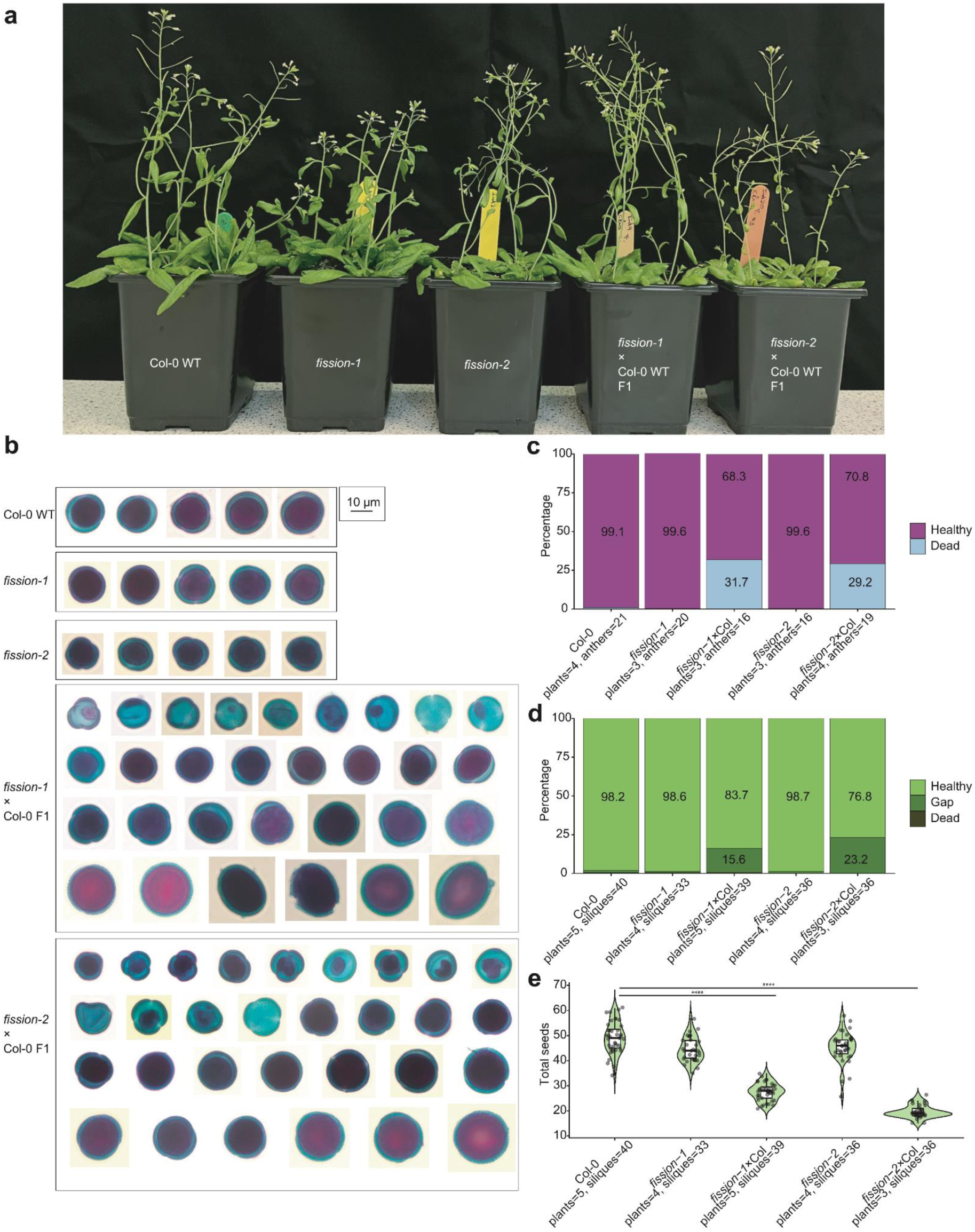
Phenotyping of wild type, fission lines, and backcrossed F_1_ hybrids. **a.** Photographs of seven to eight-week-old Arabidopsis plants. From left to right: Col-0 wild type, *fission-1*, *fission-2*, *fission-1* (maternal)×Col-0 (paternal) n=5×n=6 F_1_ hybrids, and *fission-2* (maternal)×Col-0 (paternal) n=5×n=6 F_1_ hybrids. No vegetative growth phenotype was observed in any genotype compared to wild type. **b.** Representative brightfield images of Alexander-stained pollen grains from wild type, self-fertilized fission lines, and backcross progeny of fission lines crossed to wild type (n=5×n=6). In this assay, viable pollen stains magenta/purple, whereas non-viable or aborted pollen appear blue-green. Abnormally shaped and non-viable pollen grains are observed specifically in n=5×n=6 backcross progeny, but not in wild type, or selfed fission lines. **c.** Quantification of the proportion of healthy and dead pollen grains based on Alexander staining. Only F_1_ n=5×n=6 hybrids show elevated dead pollen frequencies (31% for *fission-1*×Col-0, and 29% for *fission-2*×Col-0). **d.** Quantification of healthy, dead, and aborted (gaps) seeds per silique. Aborted ovules (gaps) are observed specifically in n=5×n=6 hybrids, ranging from 15% (*fission-1*×Col-0) to 23% (fission-2×Col-0). **e.** Quantification of total seed number per silique. F_1_ n=5×n=6 hybrids show a significant reduction in seed set compared to Col-0 wild type (Wilcoxon rank-sum test; *fission-1*×Col-0 vs Col-0 *P*=2.23×10⁻¹⁴; *fission-2*×Col-0 vs Col-0 *P*=6.28×10⁻¹⁴; ****=*P*<0.0001).

**Extended Data Figure 12.**
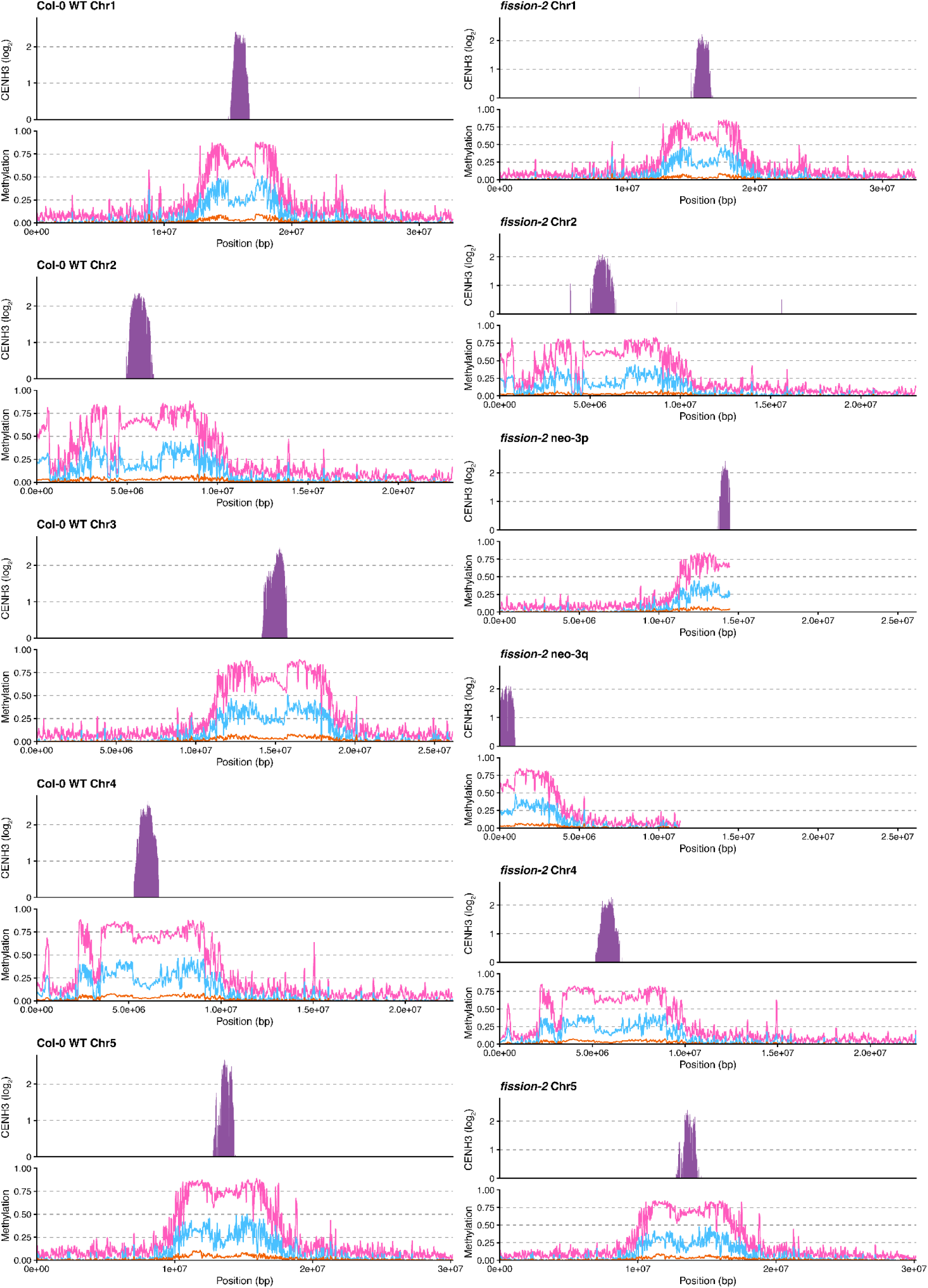
CENH3 enrichment and DNA methylation profiles across wild type and neo-chromosome centromeres. Genome-wide profiles are shown in 10 kb windows across the five chromosomes of wild type (Col-0, left) and the six chromosomes of the *fission-2* line (right). Purple histograms represent CENH3 ChIP-seq enrichment (log_2_ ChIP/input), and line plots indicate DNA methylation levels in CG (pink), CHG (blue), and CHH (yellow) sequence contexts. In the *fission-2* genome, the neo-centromeric arrays on chromosome arms neo-3p and neo-3q recruit CENH3 to levels comparable in amplitude and domain width to the other centromeres. Consistent with canonical Arabidopsis centromeres, both endogenous and neo-centromeres display a characteristic context-specific DNA methylation pattern of dense CG methylation across the centromeric satellite arrays, with relatively reduced non-CG methylation within the satellite arrays compared to the surrounding pericentromeric regions. Genomic coordinates are indicated in base pairs (bp).

**Extended Data Figure 13.**
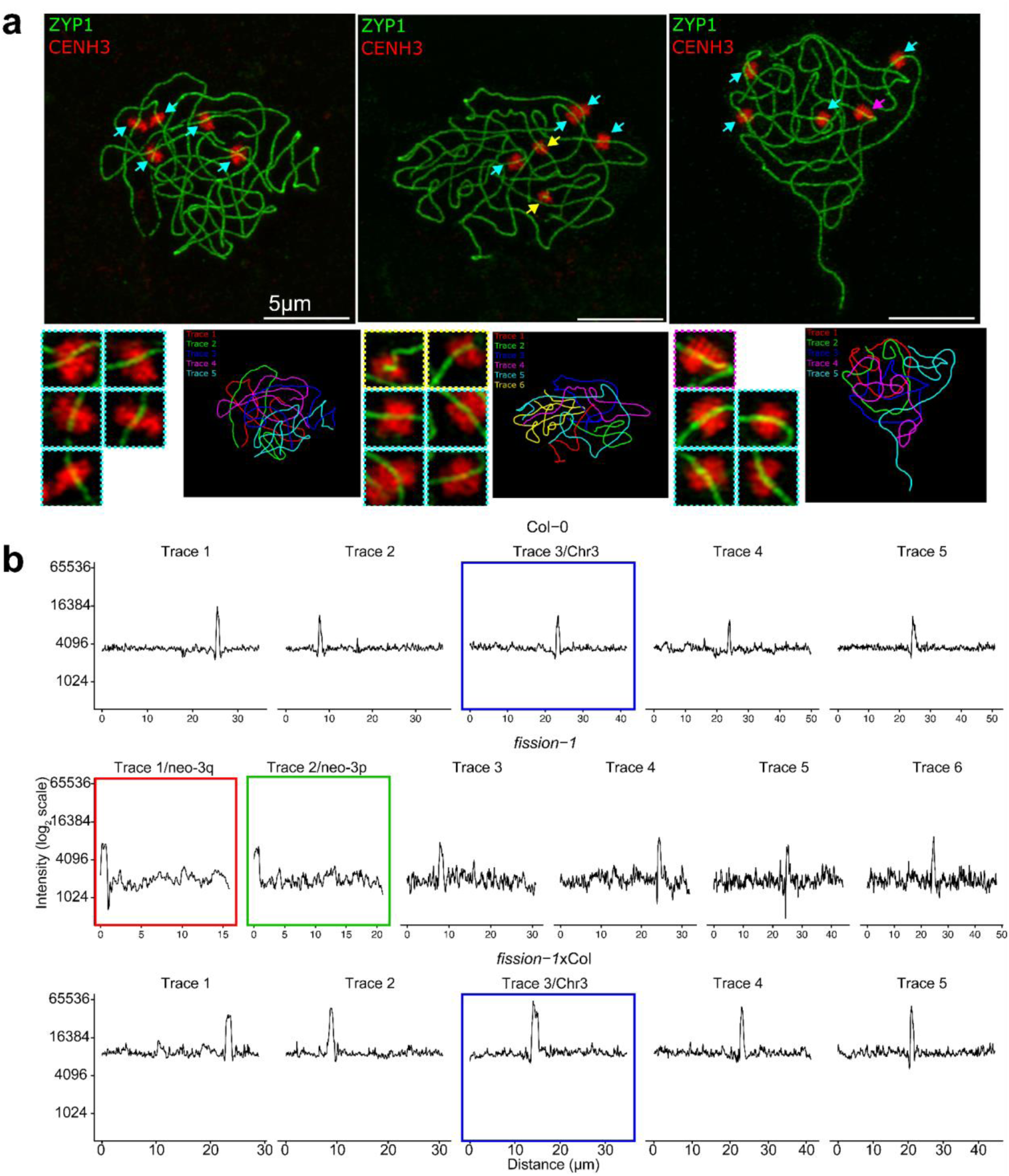
CENH3 localization on chromosome traces in Col-0, *fission-1* and *fission-1*×Col-0. **a.** Representative pachytene nuclei from Col-0, *fission-1* and *fission-1*×Col-0 immunostained for ZYP1 (green) and CENH3 (red). ZYP1 marks the synaptonemal complex and CENH3 marks centromeric chromatin. Arrowheads indicate CENH3 foci associated with individual chromosome axes. In *fission-1* the yellow arrowheads denote the neo-chromosomes while in *fission-1*×Col-0 the trivalent is highlighted by a magenta arrow. Scale bar=5 µm. Magnified views of CENH3 signals and corresponding chromosome-axis traces from the nuclei shown are also shown. Individual chromosome traces are numbered and color-coded. **b.** CENH3 fluorescence intensity profiles measured along individual chromosome traces. Profiles show CENH3 intensity as a function of distance along each chromosome axis. Traces are shown separately for Col-0, *fission-1,* and *fission-1*×Col-0. Chromosome three, or the neo-chromosomes neo-3p and neo-3q traces are indicated by colored outlines.

**Extended Data Figure 14.**
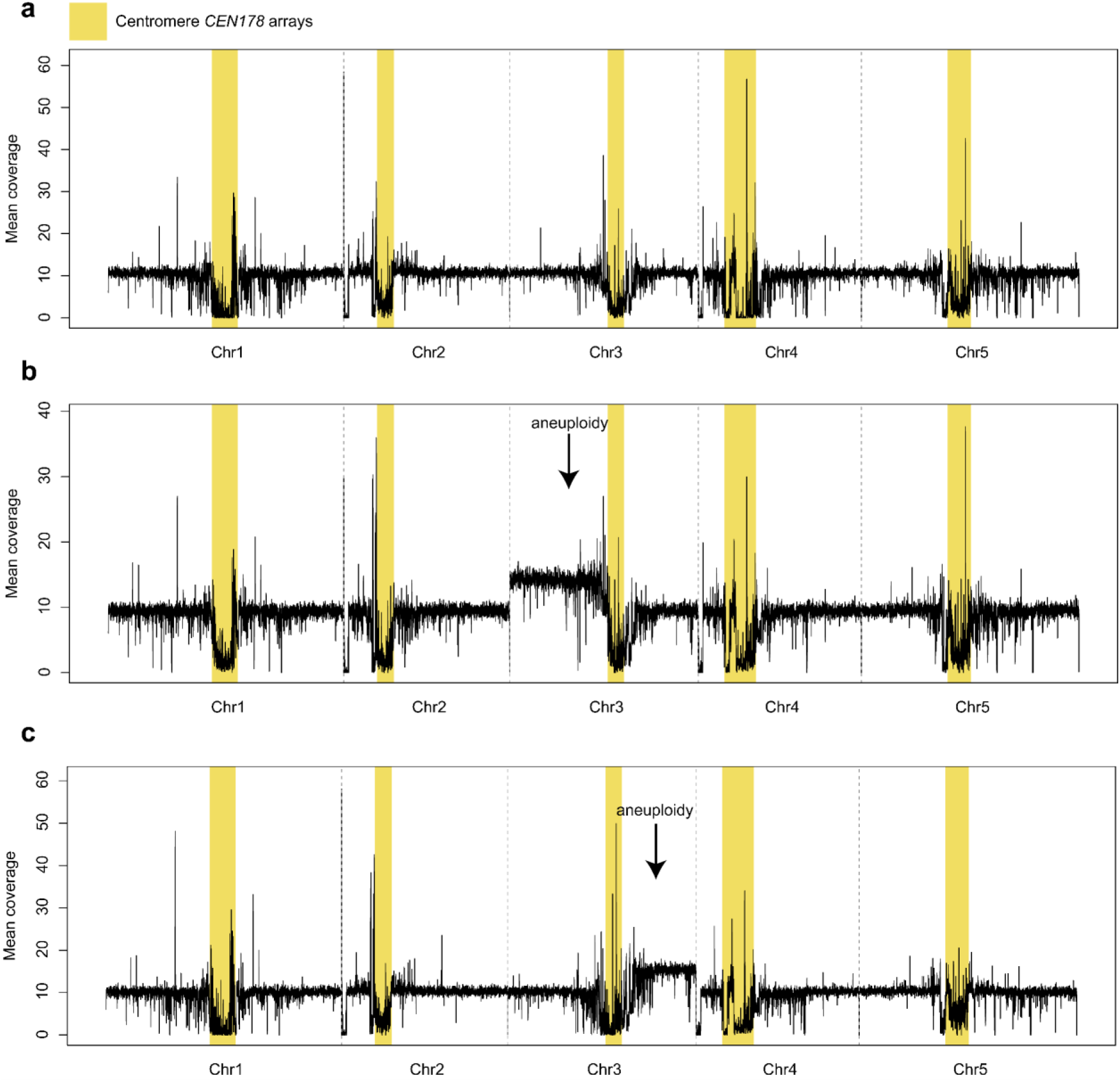
Aneuploidy of the neo-chromosomes segregating in the F_2_ generation from a hybrid cross (n=5×n=6). **a.** Following sequencing and alignment of reads from 96 F_2_ lines derived from selfed *fission-2*×Ler-0 F_1_ plants, 83 were found to be euploid. A representative example is shown. **b.** A representative of three lines that showed evidence of neo-3p triploidy. **c.** A representative of ten lines that showed evidence of neo-3q triploidy. Yellow bars highlight centromere satellite array positions, and black traces show mean DNA sequencing coverage in 10 kb windows.

**Extended Data Figure 15.**
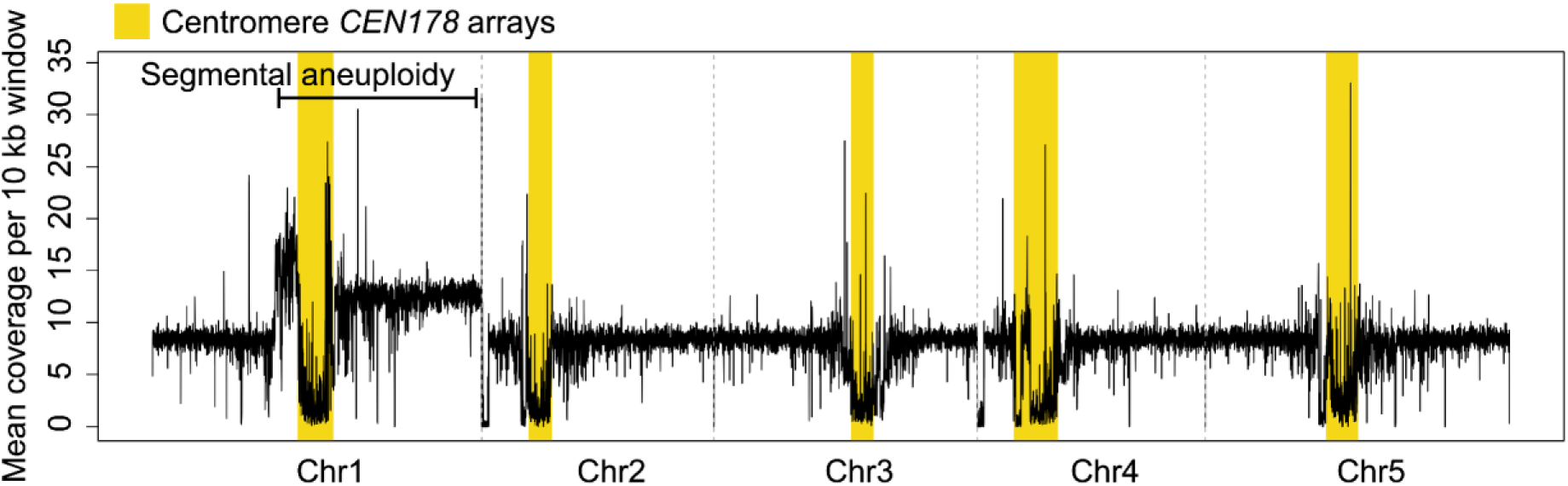
Segmental aneuploidy observed in a single individual of Col-0×Ler control F_2_ population. Shown is median DNA sequencing coverage in 10 kb from one individual from the control F_2_ population of Col-0×Ler-0 in which we observed segmental aneuploidy (trisomy) of Chr1, including the p-arm pericentromere, centromere array and q-arm of chromosome one. Yellow bars highlight centromere satellite arrays, and black traces show mean DNA sequencing coverage in 10 kb windows.

**Extended Data Figure 16.**
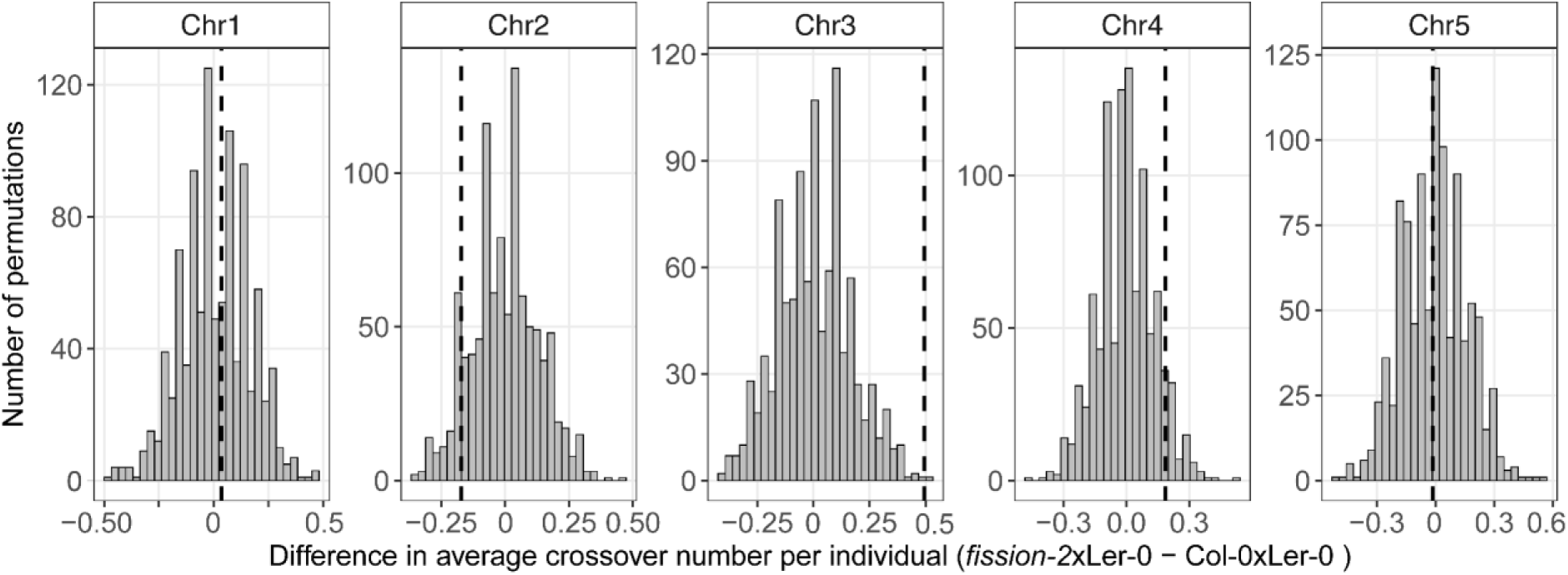
Permutation tests for crossover differences by chromosome between fission hybrids and wild type. The dashed vertical black line shows the observed difference in mean crossover number per individual between the *fission-2*×Ler-0 and control Col-0×Ler-0 F_2_ populations, for each chromosome. Grey histograms show the distribution of differences obtained from 1,000 random permutations in which individuals were randomly chosen between the two populations. For each chromosome, differences were calculated from the difference of *fission-2*×Ler-0 minus Col-0×Ler-0. *P*-values were 0.462, 0.919, 0.001, 0.102, and 0.552 for chromosomes one to five, respectively, with only chromosome three being significantly different.

**Extended Data Figure 17.**
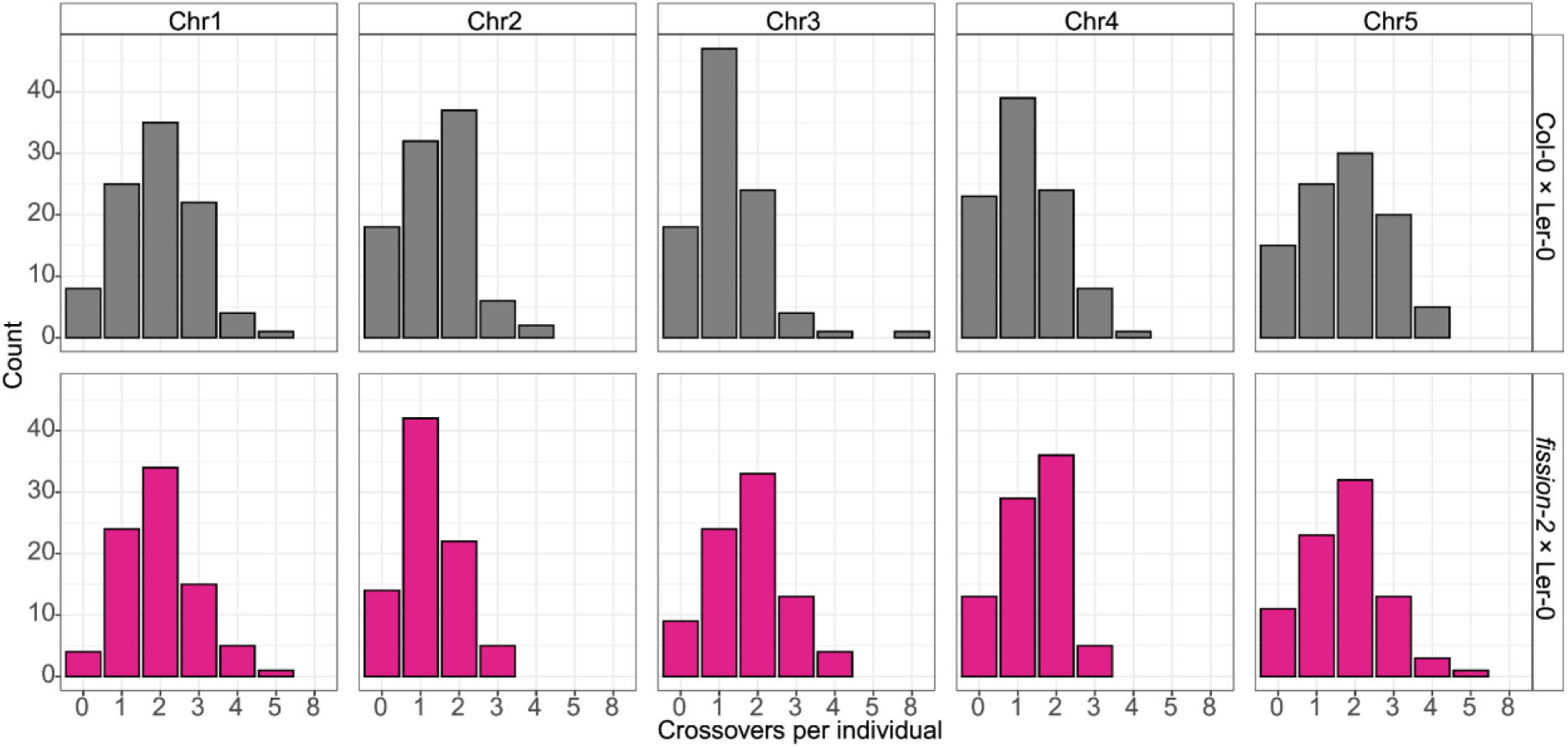
Distribution of crossovers per chromosome in wild type and n=5×n=6 fission hybrids. Barplots showing the number of crossovers per individual for each chromosome (Chr1–Chr5) in Col-0×Ler-0 control F_2_ populations (grey, top), and *fission-2*×Ler-0 n=5×n=6 hybrid F_2_ populations (pink, bottom). Across all chromosomes, crossover counts are centered around one to three events per chromosome. In chromosome three in the n=5×n=6 hybrid F_2_ population the median number of crossovers is two, compared to one in the control population, consistent with each neo-chromosome independently experiencing one obligate crossover.

**Extended Data Figure 18.**
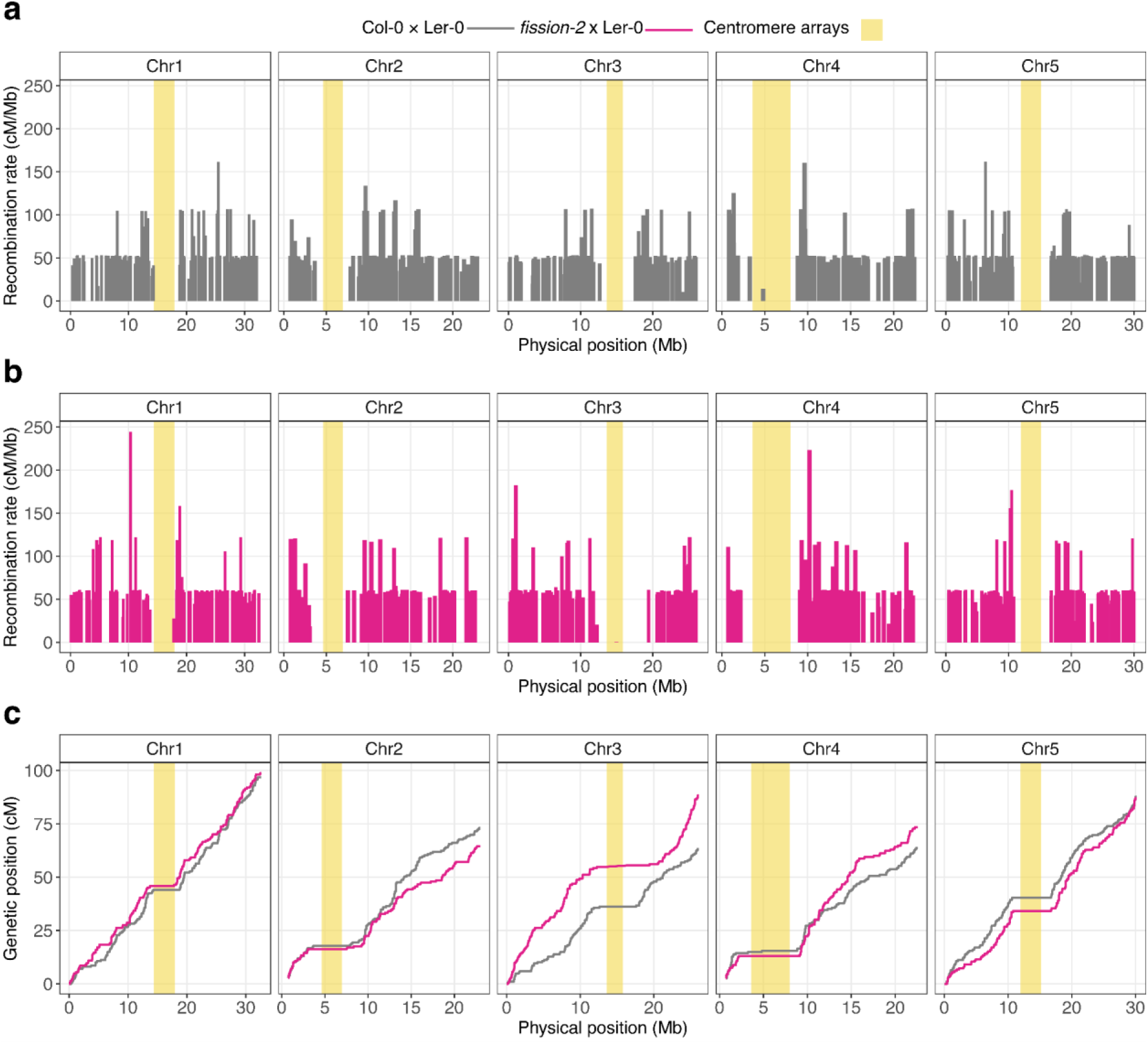
Meiotic crossover recombination landscapes in hybrid n=5×n=6 fission crosses compared with wild type controls. **a.** Crossover recombination rate across chromosomes one to five for the control Col-0×Ler-0 F_2_ population (grey). Histograms show the distribution of crossover rates along each chromosome. **b.** Crossover rate profiles across chromosomes one to five for the hybrid *fission-2*×Ler-0 n=5×n=6 F_2_ population (pink). In the *fission-2*×Ler-0 hybrid, chromosome three shows a distal redistribution of crossovers toward telomeric regions, and an expanded zone of crossover suppression in centromere-proximal regions, relative to the control. **c.** Marey maps relating physical distance (Mb) to genetic distance (cM) across chromosomes one to five in the control Col-0×Ler-0 F_2_ population (grey), and the hybrid *fission-2*×Ler-0 n=5×n=6 F_2_ population (pink). Yellow shading marks the *CEN178* arrays for each chromosome.

**Extended Data Figure 19.**
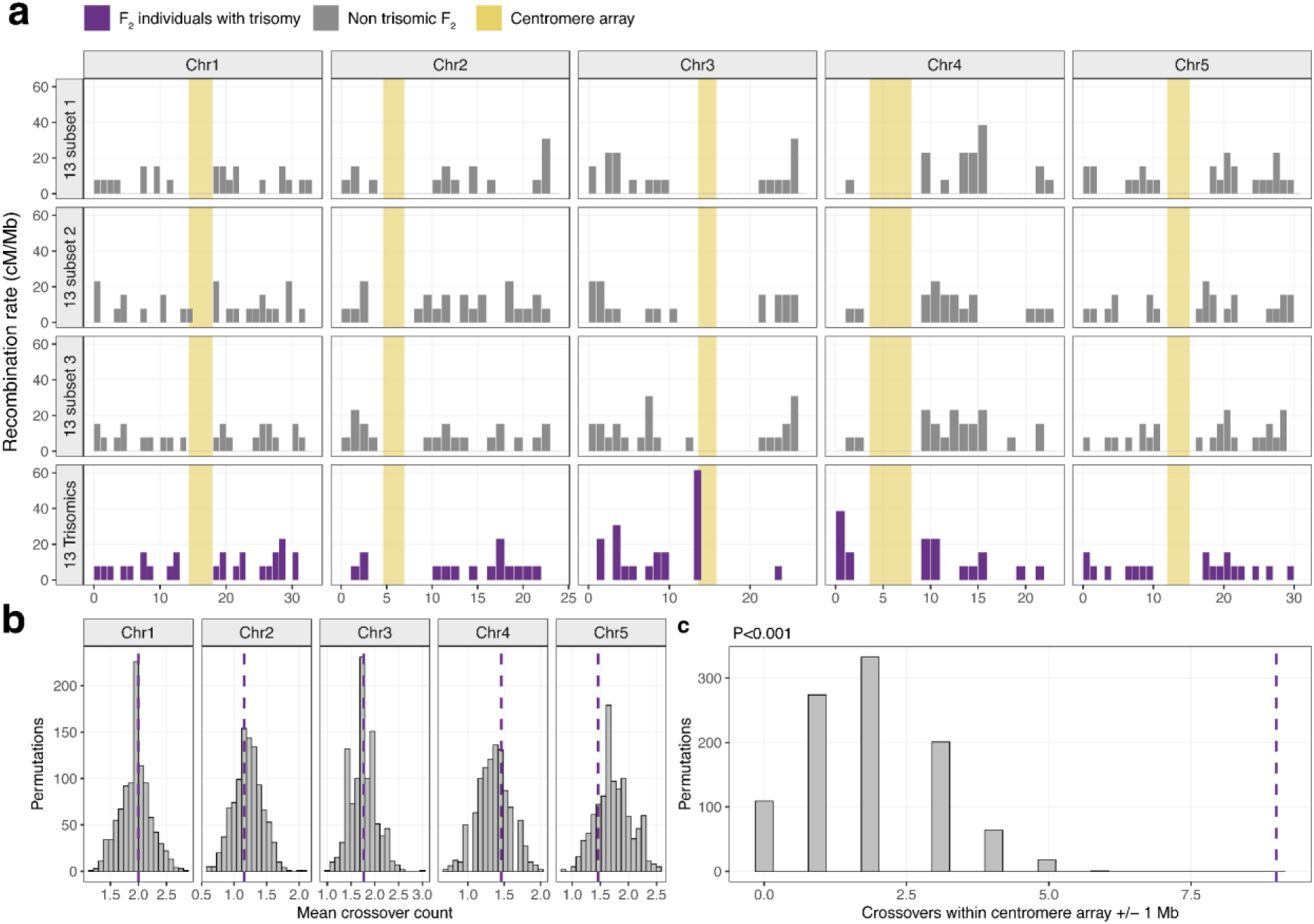
Crossover profiles and permutation analyses in trisomic F_2_ individuals. **a.** Recombination rate profiles across all five chromosomes for three random subsets of 13 non-trisomic F_2_ individuals and the 13 trisomic F_2_ individuals. Bars show recombination rate in 1-Mb windows, expressed as cM/Mb. Grey bars indicate non-trisomic F2 subsets and purple bars indicate trisomic F_2_ individuals. Yellow shading marks the centromere array on each chromosome. Crossover intervals spanning a centromere array were retained but plotted at the nearest centromere-array edge. **b.** Null distributions from 1,000 permutations of 13 non-trisomic F_2_ individuals showing the mean crossover count per chromosome. Purple dashed lines indicate the observed mean crossover count in the 13 trisomic F_2_ individuals. No significant increase of the total number of crossovers in the trisomic individuals was observed. **c.** Permutation test for centromere-proximal (i.e. crossovers falling within the centromere array ±1 Mb). Grey bars show the null distribution from 1,000 permuted sets of 13 non-trisomic F_2_ individuals, and the purple dashed line shows the observed trisomic value. The observed trisomic enrichment exceeded all permutations, giving P < 0.001. While trisomic individuals did not show a significant increase in total crossover number, crossover positioning was altered, with enrichment of chromosome 3 centromere-proximal crossovers.

**Extended Data Figure 20.**
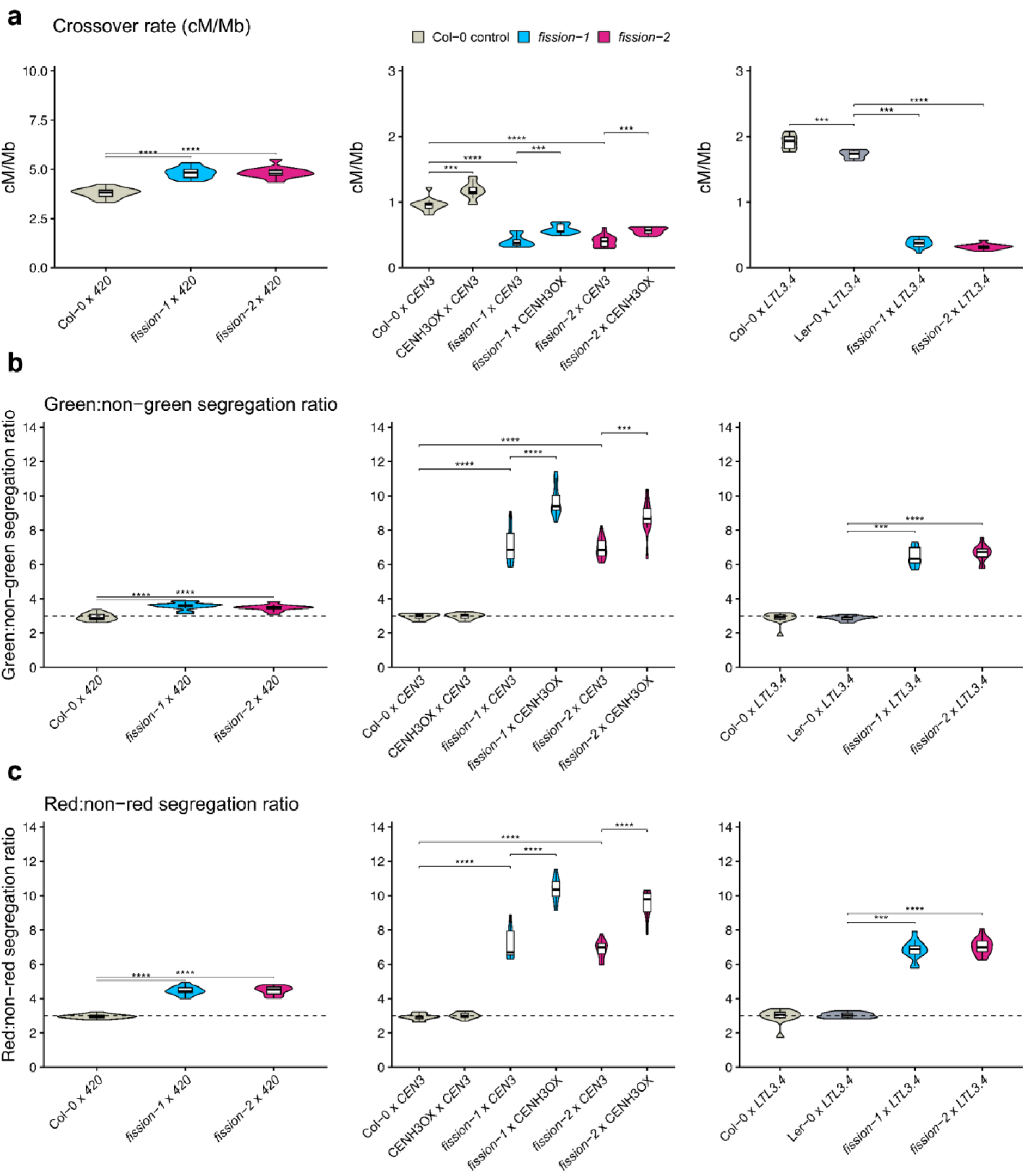
Meiotic crossover frequency and segregation distortion in wild type and fission hybrids assayed using fluorescently tagged lines. **a.** Crossover rate (cM/Mb) across the *420*, *CEN3*, and *LTL3.4* intervals in control, *fission-1*, *fission-2*, and CENH3-overexpression backgrounds. **b.** and **c.** Green:non-green (G:nonG) (**b**) and red:non-red (R:nonR) (**c**) segregation ratios across the same intervals and genotypes. Each point represents an independent F_2_ population. Violin plots show the ratio distribution across replicate plants. Overlaid box plots indicate the median and interquartile range. The dashed line indicates the expected Mendelian 3:1 segregation ratio. Statistical comparisons used two-sided Wilcoxon rank-sum tests. \*\*\**P* < 0.001; \*\*\*\**P* < 0.0001.

**Extended Data Figure 21.**
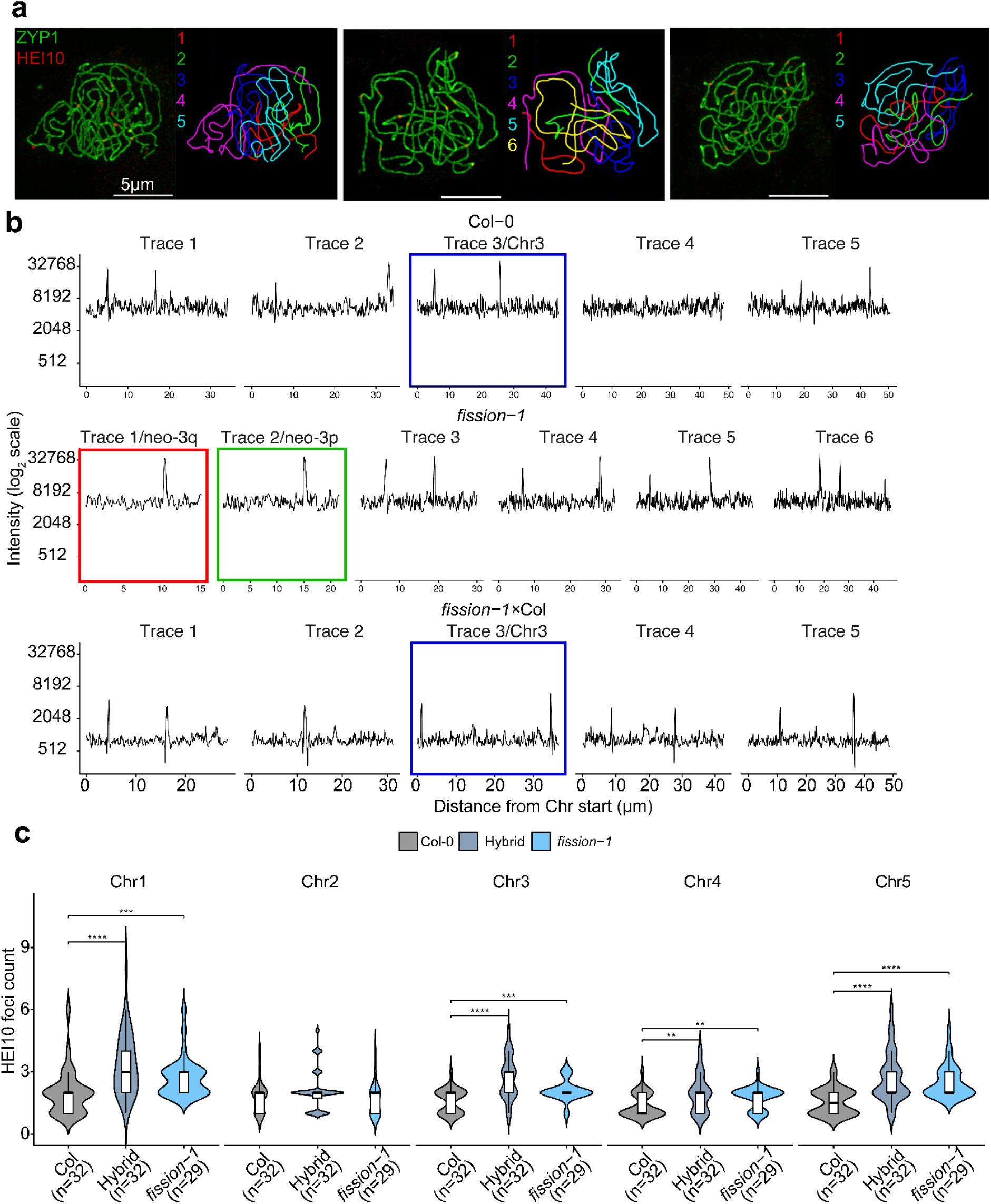
HEI10 patterning on individual chromosome traces in Col-0, *fission-1*, and *fission-1*×Col. **a.** Pachytene nuclei from Col-0, *fission-1*, and *fission-1*×Col-0 immunostained for ZYP1 and HEI10 with traces. ZYP1 marks the synaptonemal complex and HEI10 marks crossover-associated foci. Individual chromosome traces are numbered and color-coded. Chromosomes were discerned based on tracing of the synapse chromosomes. Scale bar=5 µm. **b.** HEI10 fluorescence intensity profiles measured along individual chromosome traces from the nuclei shown in A. Profiles show HEI10 intensity as a function of distance along each traced chromosome. For *fission-1*, the two neo-chromosome traces corresponding to the split chromosome three configuration are shown separately as neo-q and neo-p. Chromosome three or the corresponding neo-chromosome traces are indicated by colored outlines. **c.** Quantification of HEI10 foci number per chromosome for chromosomes one to five in Col-0, *fission-1*, and *fission-1*×Col-0. Violin plots show the distribution of foci counts, with boxplots indicating the median and interquartile range. Statistical comparisons were performed between Col-0 and each fission genotype using two-sided Wilcoxon rank-sum tests. Significance levels are indicated as as asterisks *P*<0.001 (***) *P*<0.0001 (****).

**Extended Data Figure 22.**
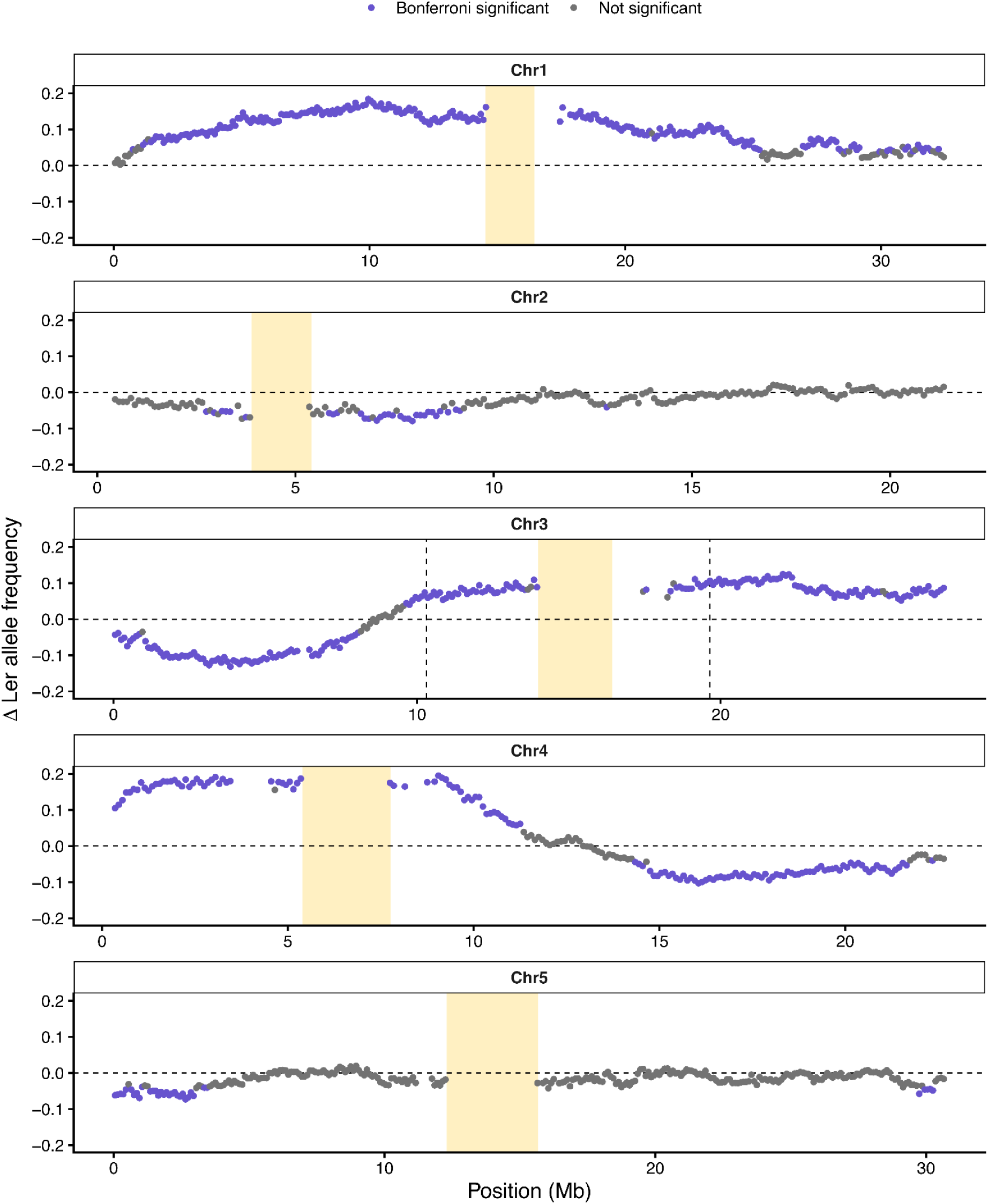
Genome-wide segregation distortion in the *fission-2*×Ler-0 F_2_ population compared to controls. Ler-0 allele frequency differences between the *fission-2*×Ler-0 and Col-0×Ler-0 F_2_ populations were calculated in non-overlapping 100 kb windows across all five Arabidopsis chromosomes. Each point represents one window and shows the change in Ler-0 allele frequency, calculated as *fission-2*×Ler-0 allele frequency minus Col-0×Ler-0 frequency. Purple points indicate windows that were significantly different after Bonferroni correction; grey points are not significant. Significance was tested using Fisher exact test. Yellow shading marks Ler-0 centromeric *CEN178* arrays.

**Supplementary Table 1.**
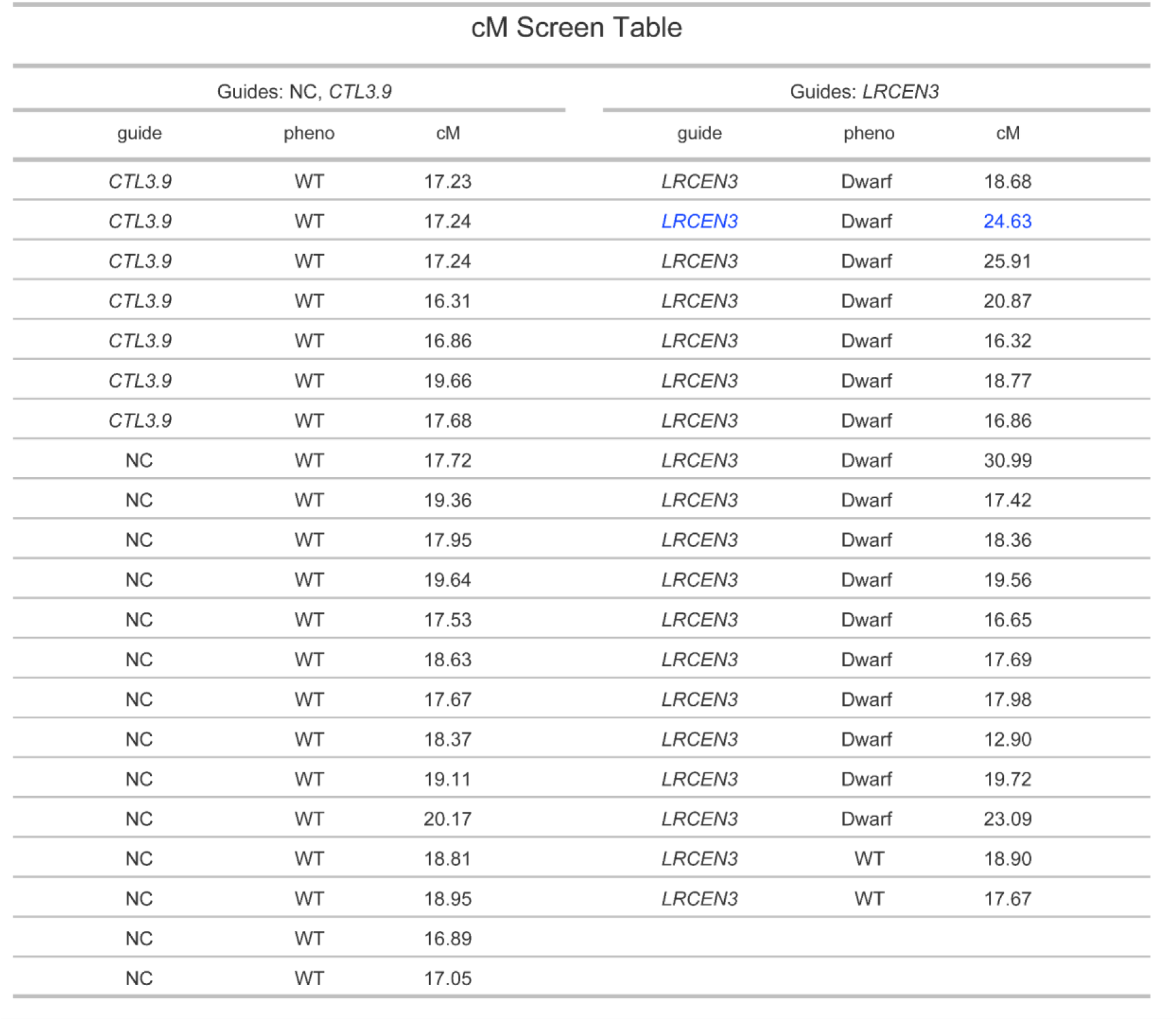
Genetic distances of *CTL3.9* controls and *CRISPR–CEN3* transgenic lines. The table summarizes crossover recombination distances (centiMorgans, cM) measured in hemizygous *CTL3.9* plants. Two controls are included: untransformed *CTL3.9* plants and a no-guide Cas9 negative control (NC). The CRISPR guide *LRCEN3* corresponds to T₁ lines expressing Cas9 with the centromere three targeting guide RNA. The sample highlighted in blue was advanced to the T₂ generation for further analysis.

**Supplementary Table 2.** Summary of contig-level assembly of neo-chromosome lines. Contig-level assembly statistics. The N50 and N90 values represent the contig lengths at which 50% and 90% of the total assembly length are contained in contigs of that size or longer, respectively. The L50 and L90 values indicate the minimum number of contigs required to reach 50% and 90% of the total assembly length. The auN (area under the Nx curve) provides an integrated, genome-wide measure of assembly contiguity across all Nx thresholds. Statistics were generated using RagTag (*2*).

| Sample | n (contigs) | Total bp | Min (bp) | Max (bp) | auN (bp) | N50 (bp) | N90 (bp) | L50 | L90 |
| --- | --- | --- | --- | --- | --- | --- | --- | --- | --- |
| *fission*<br>-1 | 108 | 154,757,209 | 28,671 | 32,635,998 | 21,894,798 | 22,865,054 | 6,104,200 | 3 | 7 |
| *fission*<br>-2 | 150 | 153,727,927 | 18,841 | 32,636,472 | 22,052,276 | 23,072,733 | 6,096,738 | 3 | 7 |

**Supplementary Table 3.**
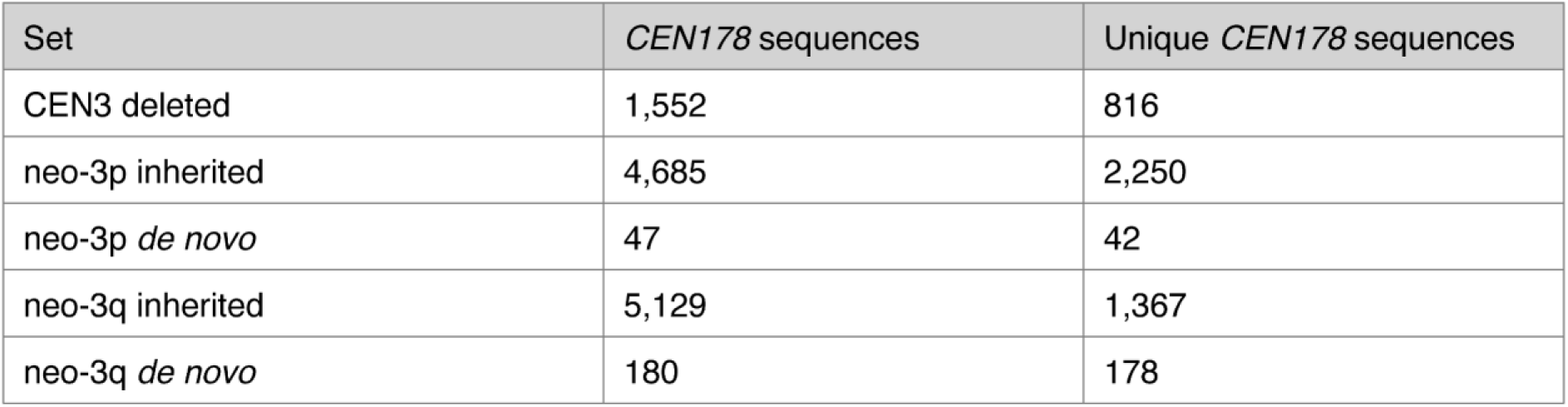
Summary of *CEN178* changes to the neo-3p and neo-3q neo-centromere arrays. Distribution of *CEN178* sequences across deleted, inherited, and *de novo* categories in the neo-3p and neo-3q neo-centromere arrays. Total counts and unique (non-redundant) sequences after deduplication are shown for each set.

## Materials and Methods

### Design of *CRISPR-CEN3* guide RNA sequence

We designed centromere-specific guide RNAs (gRNA) for *A. thaliana* Col-0 using the Col-CEN genome assembly^28^. All potential Cas9 guides were identified using a string search for the ‘NGG’ protospacer adjacent motif (PAM). For each candidate guide sequence, we calculated the number of target sites within the *CEN178* repeat arrays, in comparison to those located outside of the repeat arrays. Guides with at least 100 centromere-specific target sites, and no off-targets elsewhere in the genome were retained for further analysis. From the filtered guide RNA list, we selected a sequence that targets centromere three with the highest target count, and a relatively even distribution within the *CEN178* arrays. The sequence of the *CRISPR-LRCEN3* guide RNA is 5’-AGGCTTACAAGATTGGGTTG-3’, which has 154 predicted targets with the NGG PAM on centromere three, and no off-targets using a perfect match. Centromere three was prioritized because of the availability of flanking fluorescent markers for the *CTL3.9* traffic line^30^, facilitating downstream analyses of screening for structural changes by looking for deviations from expectations of recombination between the fluorescent marks.

### Cloning of the *CRISPR-LRCEN3* constructs

The selected *CRISPR-LRCEN3* guide RNA sequence was introduced into the plasmid vector pYAO-Cas9, which carries kanamycin and hygromycin resistance cassettes for selection of transformed bacterial colonies and transformed plants, respectively^29^. The pYAO promoter drives efficient Cas9 expression in the early embryo and meristems, in addition to being expressed in pollen, the embryo sac, embryo, and endosperm^29,48^. To introduce the *CRISPR-LRCEN3* guide RNA sequence, DNA constructs of *ATU6-26-sgRNA* were amplified using high-fidelity Phusion-PCR polymerase with overlapping 15 bp ends. The guide RNA sequence is introduced immediately downstream of the *ATU6-26* promoter. The restriction enzyme *Spe*I was used to linearise the pYAO-Cas9 plasmid. The *SpeI* cut site is 31 bp upstream of the pYAO promoter. The *CRISPR-LRCEN3* DNA constructs were inserted into the pYAO-Cas9 vector using In-Fusion assembly and a 3:1 insert:vector ratio. The resulting plasmid was cloned into DH5α E. coli cells by heat-shock. Whole plasmid sequencing from positive cells was performed by Plasmidsaurus using Oxford Nanopore Technology with custom analysis and annotation. To determine whether the observed dwarf phenotype in transformed plants was due to Cas9 endonuclease activity, a catalytically inactive Cas9 (dCas9) was used. The *CRISPR-LRCEN3* guide RNA was cloned into the *UBQ10*:dCas9 expression vector^49^, in which we removed the sequence encoding the TET1 catalytic domain in the vector using the restriction enzyme *Bsi*WI. Transformation of the *UBQ10*:dCas9-*LRCEN3* vector enabled targeted recruitment of dCas9 to the centromere three target region without introducing DNA double-strand breaks. Plant transformation was carried out via Agrobacterium floral dip method using the strain GV3101 and transformed T_1_ plants were selected on medium containing hygromycin.

### Plant transformation, growth conditions, and pedigree

For transformation, we used hemizygous *CTL3.9* plants that were generated from a cross between wild type Col-0 and the homozygous traffic line *CTL3.9*^30^. *CTL3.9* RG/++ hemizygous plants were transformed using the floral dip method and Agrobacterium tumefaciens strain GV3101^50^. Plants were grown in controlled environment chambers at 20°C under long day conditions (16/8 hours light/dark photoperiods), with 60% humidity, and 150 µmol m^-2^ second^-1^ light intensity. The *CRISPR-LRCEN3* construct was transformed into hemizygous *CTL3.9* RG/++ via floral dipping (T_0_ generation). T_1_ seeds were grown on selection plates. Selected T_1_ plants were allowed to set seed, and the seed was scored for *CTL3.9* genetic distance (cM). In the T_2_ generation, non-fluorescent seeds were selected to remove the FTL T-DNA markers. Plants with a dwarf developmental phenotype were identified and a single T_2_ line with the dwarf phenotype was carried forward to the T_3_ generation. Twenty sibling T_3_ individuals were screened for karyotypic alterations using FISH and for the absence/presence of the Cas9 construct. From the twenty T_3_ siblings, seven non-dwarf plants that did not contain the Cas9 construct but did have a karyotype change were retained. Two of the seven T_3_ individuals with a novel karyotype of 2n=12 were carried forward to the T_4_ generation (T_4_-4, and T_4_-20). These individual lines were respectively renamed *fission-1* and *fission-2* and genomes were sequenced and assembled from these lines using Oxford Nanopore Technologies (ONT) sequencing. T_5_ progeny from these fission lines were used for RNA-seq, ChIP-seq, and crossing experiments.

### Measuring *CTL3.9* genetic distance using fluorescence microscopy

The *CTL3.9* line contains T-DNAs expressing red or green fluorescent proteins driven by the *NapA* promoter, which are integrated at positions 9,748,689 to 18,661,551 on chromosome three that flank the centromere^30,31^. Seeds from hemizygous *CTL3.9* RG/++ plants were observed using a fluorescent stereomicroscope (Leica M165FC). Seeds were imaged under three conditions: transmitted light illumination, a UV light with a dsRed filter, and a UV light with a GFP filter. Image analysis was performed using CellProfiler software (version 2.1.1)^51^. Seed boundaries were delineated, and fluorescence intensity values for dsRed and GFP were recorded for each seed object. The genetic distances in centimorgans (cM) for each hemizygous *CTL3.9* RG/++ plant were calculated using the formula:

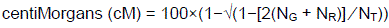

where N_G_, N_R_ and N_T_ represent the total green, total red, and overall total seed number, respectively. T_1_ plants exhibiting outlier genetic distances and a dwarf phenotype were selected for advancement to the T_2_ generation. To compare variance between groups we used the Fligner-Killeen test. To remove fluorescent markers, plants were screened under the fluorescent stereomicroscope and seeds that were homozygous null for the fluorescent FTL markers were selected.

### Cytogenetic analysis of centromere-edited lines

Both mitotic and meiotic chromosome spreads were prepared from anther tissue collected at appropriate developmental stages. The material was fixed in ethanol:acetic acid (3:1) and stored in 70% ethanol until use. Chromosome preparations were obtained according to the protocol described^52^. For chromosome identification, Arabidopsis chromosome-specific BAC clones corresponding to the left (p) and right (q) arms of chromosome three were used. The p arm was identified using the BAC clones T4P13, T13O15, F1C9, F16B3, T21P5, T11I18, F7O18, F22F7, F2O10, and F24P17, whereas the q arm was identified using the BAC clones F9D24, F17J16, F24G16, T8B10, T20K12, F21F14, F26K9, and F16M2. Additional probes included the *CEN178* centromeric satellite repeat^28^, the Arabidopsis-type telomeric repeat (TTTAGGG)n^53^, and the 35S rDNA probe represented by the Arabidopsis BAC clone T15P10. All DNA probes were labeled with biotin-dUTP, digoxigenin-dUTP, or Cy3-dUTP by nick translation. Labeled probes were pooled as required, ethanol-precipitated, and applied to pepsin-treated and ethanol-dehydrated slides containing suitable chromosome spreads. Probe and chromosomal DNA were co-denatured at 80°C for 2 minutes, followed by hybridization at 37 °C for 12 hours. Hybridized probes were detected by immunofluorescence as described by Mandáková and Lysak (2016). Biotin-labeled probes were detected using avidin-Texas Red (Vector Laboratories), and digoxigenin-labeled probes were detected using mouse anti-digoxigenin antibodies (Jackson ImmunoResearch) followed by goat anti-mouse Alexa Fluor 488 or Alexa Fluor 647 (Invitrogen). Chromosomes were counterstained with DAPI (2 µg/mL) in Vectashield (Vector Laboratories). Fluorescence signals were analyzed using a Zeiss AxioImager epifluorescence microscope (Carl Zeiss) equipped with a CoolCube camera (MetaSystems). Images were acquired separately for each fluorochrome using Isis software (MetaSystems) with appropriate excitation and emission filters (AHF Analysentechnik). Monochromatic images were pseudocolored, merged, and processed using Photoshop CS (Adobe Systems). Chromosome lengths were measured using ImageJ (National Institutes of Health).

Three-dimensional immuno-FISH was performed on mitotic chromosomes of tapetal cells isolated from anthers. Anthers at appropriate developmental stages were fixed in freshly prepared 4% paraformaldehyde in 1×PBS for 30 minutes at room temperature, washed three times in 1×PBS, and embedded in polyacrylamide to preserve the native three-dimensional nuclear architecture, as described previously. After polymerization, samples were subjected to partial enzymatic digestion of cell walls to facilitate antibody penetration. Following digestion, preparations were permeabilized in 0.5% Triton X-100 in 1×PBS for 10 minutes and blocked in 3% BSA in 1×PBS for 30 minutes at room temperature. Immunolabeling was performed using a primary antibody against the centromere-specific histone variant CENH3^54^, diluted in blocking solution and incubated overnight at 4°C. After washing in 1×PBS, samples were incubated with fluorophore-conjugated secondary antibodies for 2 hours at room temperature in the dark. Following immunodetection, antibody complexes were post-fixed in 4% paraformaldehyde for 10 minutes to stabilize the immunosignals. Subsequently, preparations were subjected to FISH following the procedure described above, using the same Arabidopsis chromosome-specific BAC clones spanning the p and q arms of chromosome three. After FISH, DNA was counterstained with DAPI and samples were mounted in antifade medium. Three-dimensional image stacks were acquired using a Zeiss AxioImager epifluorescence microscope equipped with an ApoTome module. Optical sections covering the entire nuclear volume were collected and used for three-dimensional reconstruction, signal rendering, and spatial analyses in IMARIS software (Oxford Instruments). Representative 3D projections and rotational movies were generated from reconstructed mitotic chromosomes.

For 3D-SIM imaging of late-pachytene nuclei, sample preparation was performed as described^55^. In brief, anthers of the correct stage were dissected from buds and macerated using a brass rod on a No. 1.5H coverslip (Marienfeld) in 10 µl digestion medium (0.4% cytohelicase, 1.5% sucrose, 1% polyvinylpyrolidone in sterile water) for 1 minute. Coverslips were then incubated in a moist chamber at 37 °C for 3 minutes before adding 10 µl of 1.8% lipsol solution, followed by 20 µl 4% paraformaldehyde (pH 8). Coverslips were dried and then blocked in 0.3% bovine serum albumin in 1× phosphate-buffered saline (PBS) solution. Coverslips were incubated with primary antibody at 4 °C overnight and secondary antibody at 37 °C for 2 hours. In between antibody incubations, coverslips were washed 3 times for 5 minutes in 1×PBS plus 0.1% Triton X-100. Coverslips were mounted on a slide in 7 µl Vectashield with DAPI. The following antibodies were used: anti-HEI10 (rabbit, 1:500), anti-CENH3 (rabbit, 1:500), anti-ZYP1 (rat,1:500), anti-ASY1 (guinea-pig, 1:500), anti-rat Alexa Fluor 488 (AB_2896331, goat, 1:200), anti-rabbit Alexa Fluor 555 (AB_2535851, goat, 1:200), and anti-guinea-pig Alexa Fluor 647 (AB_2535867, goat, 1:200). Three-dimensional structured illumination microscopy (3D-SIM) was performed using a Zeiss Elyra7 microscope equipped with 2xPCO edge 4.2 sCMOS cameras, a Plan-Apochromat ×63, NA 1.40 oil objective, 405, 488, 561 and 640 nm lasers, and ZEN 3.13 acquisition software. Slides were imaged at 30°C in 3D SIM mode with thirteen phases according to the microscope manufacturer’s instructions, using an immersion oil with a refractive index of 1.518 optimised for imaging at this temperature. Z-stacks were captured at an interval size of 0.1 μm, using consistent microscope laser power and camera gain values for each image. For HEI10 foci detection in late-pachytene nuclei, individual bivalents were traced using the ZYP1 image channel and the SNT plugin in Fiji^56^, with analysis being restricted to nuclei in which all chromosomes were fully synapsed. Specific chromosomes were identified based on SC length, with chromosome three being identified as the bivalent/trivalent with the third longest SC length in the Col and *fission-1*xCol-0 data, and with neo-3q and neo-3p being identified as the shortest and second-shortest bivalents in the *fission-1* data. HEI10 foci identification along each bivalent was then carried out using the FociMapper analysis tool^57^, using the following parameter values: xy_res: 0.0313 µm, z_res: 0.1 µm, sphere_radius: 0.2 µm, peak_threshold: 0.25, screening_distance: 10, threshold_type: per-cell.

Pollen fertility was assessed following the protocol described^58^. Briefly, anthers from mature flowers were isolated and mounted on microscope slides. A few drops of Alexander stain were added, after which the samples were covered with a coverslip and gently squashed by hand. After 15 minutes, the slides were examined under a light microscope.

### Oxford Nanopore sequencing and genome assembly of fission lines

Three-week-old seedlings of the fission lines were grown under standard conditions on half-strength Murashige and Skoog (½ MS) medium supplemented with 1% (w/v) sucrose. Approximately 2 grams of fresh tissue was harvested, flash-frozen in liquid nitrogen, and ground into a fine powder using a pre-chilled mortar and pestle. High molecular weight (HMW) genomic DNA was extracted from the ground tissue using the NucleoBond HMW DNA kit (Macherey-Nagel), following the manufacturer’s protocol optimized for plant material. To enrich long DNA fragments and remove shorter molecules the extracted DNA was processed using the Short Read Eliminator (SRE) kit from PacBio. For library preparation, barcoded adapters were ligated to the DNA using the Oxford Nanopore Native Barcoding Kit, compatible with the v14 sequencing chemistry, and R10.4 flow cells. DNA libraries were subsequently sequenced on a PromethION platform using R10.4 flow cells. Base-calling was performed using Dorado (version 0.9.5). Promethion sequencing of the fission lines produced a total of 3.46 million reads, and an average read N50 of 22.7 kb. Genomes were assembled using Hifiasm (version 0.25.0) with the option ‘--ont’ which accepts R10 ONT reads, rather than PacBio Hifi reads^59^.

### Centromere and telomere sequence analysis

Centromere tandem repeat arrays were identified using the software TRASH^60^. Telomere locations were identified using the software tidk^61^. Divergence to the consensus telomere sequence was calculated by examining the first and last 4 kb of each chromosome. The short arms of chromosomes two and four were excluded from the analysis, because the end of these chromosomes include the ribosomal DNA (rDNA) arrays, which prevent accurate telomere assembly for these chromosome arms. DNA sequences from each 4 kb chromosome end were scanned with a 7 bp sliding window corresponding to the canonical telomere repeat (5’-TTTAGGG-3’), considering all circular rotations of the motifs. We calculated the minimum Hamming distance between the local 7-mer, and the canonical rotations to quantify mismatches to the telomere repeat, ranging from 0 to greater than 3.

### CENH3 chromatin immunoprecipitation and sequencing

Approximately four grams of two-week-old seedlings were collected and immediately flash-frozen in liquid nitrogen. The frozen tissue was ground into a fine powder using a Qiagen Retsch TissueLyser and a steel grinding ball. Crosslinking, nuclei isolation and chromatin recovery were performed following the protocol described by^62^. Chromatin was sonicated using a Covaris E220 with the following parameters: power=150 W, duty factor=20%, 200 bursts per cycle, for 90 seconds. The fragmented chromatin was incubated overnight at 4°C with 50 µl of Protein A magnetic Dynabeads (Thermo Fisher), pre-bound with 5 µl of a α-CENH3 antibody^32^. Sequencing libraries were prepared using the Ovation Ultralow System V2 DNA-sequencing library preparation kit from Tecan. Pre-made libraries were sequenced at Novogene using 150-bp paired-end reads, generating 6 Gb of data for the CENH3 ChIP and 12 Gb for the input control. ChIP-seq reads were aligned to the Col-0 assembly using Minimap2^63^ (v2.26) in short-read mode^22^. Alignments were sorted and indexed with SAMtools (v 1.21). Secondary, supplementary and unmapped alignments were removed and per-base genome coverage was calculated with SAMtools. When correlating log₂(CENH3/Input) enrichment between wild type and fission lines, we retained *CEN178* sequences that had a log₂(CENH3/Input) greater than 1 in the fission line on the neo-3p array, while the control had log₂(CENH3/Input) of 0. To account for background noise a tolerance threshold of 1×10⁻⁹ was applied to *CEN178* during the analysis.

### DNA methylation analysis

Oxford Nanopore sequencing data (R10) were basecalled using Dorado (v1.3.1) with modified base calling enabled. Basecalling was performed from POD5 files using an R10.4.1 super-accuracy model trained for joint detection of 5-methylcytosine (5mC) and 5-hydroxymethylcytosine (5hmC) (dna_r10.4.1_e8.2_400bps_sup@v5.2.0_5mC_5hmC@v2). Modified base calling was conducted during basecalling, enabling simultaneous inference of nucleotide sequence and cytosine modification status. Modified base information was retained in BAM files as MM and ML tags for downstream analysis. Cytosine methylation was analyzed across CpG, CHG, and CHH sequence contexts. Downstream processing and summarization of modified base calls were performed using modkit (v0.6.0). Modkit was used to extract per-site methylation information from BAM files and to calculate methylation levels as the proportion of modified cytosines relative to the total number of callable cytosines at each genomic position. To assess methylation differences between the neo-3p centromere array and the intact array of wild-type *CEN3*, we used per-site methylation estimates generated with modkit. Methylation levels were calculated for each shared *CEN178* sequence based on its start and end coordinates, and methylation patterns between the two arrays were compared across the CG, CHG, and CHH sequence contexts.

### RNA sequencing and gene expression analysis

Plants were grown at 21°C and harvested at the stage of seven true leaves, with three replicates per genotype. Total RNA was extracted from individual leaves using TRIzol reagent and DNA was removed using TURBO DNA-free kit. For flower bud expression analysis, unopened buds were collected and flash-frozen in liquid nitrogen, with three biological replicates per genotype. For both leaves and flower buds, RNA sequencing libraries were prepared by Novogene, generating strand-specific polyA libraries. Sequencing was performed on the NovaSeq X Plus Series platform, generating paired-end 150 base pair reads with a total of 6 Gb of sequencing data per sample. Reads were aligned to the PacBio Hifi genome of Col-0^22^, using STAR (version 2.7.11), retaining only uniquely mapped reads^64^. Genes with low expression were filtered out by setting a minimum transcripts per million (TPM) threshold of 3 across at least two samples. Differentially expressed genes (DEGs) were identified using the DESeq2 package in Python^65^.

### Fission line genetic analysis

To examine meiotic recombination and segregation distortion across centromere three, we used the fluorescent tagged line (FTL) *CEN3*, which carries a red fluorescent T-DNA marker (CR880) at base pair position 12,652,136 and a green fluorescent T-DNA marker (CG811) at position 18,395,802, spanning 5.74 Mb on chromosome three in Arabidopsis^32^. Wild type Col-0 and the fission lines were crossed to *CEN3* homozygotes (RG/RG), and recombination frequency and segregation distortion were quantified by scoring inheritance of the fluorescent markers. Under Mendelian expectations, self-fertilization of the resulting F_1_ plants yields a 3:1 ratio of fluorescent to non-fluorescent progeny. To assess the effect of altered CENH3 dosage, Col-0 and the fission lines were crossed to *CEN3* in the RG/RG background carrying a CENH3 overexpression construct (*CENH3-OX*). In this line, *CENH3* is expressed from the constitutive *RPS5a* promoter in actively dividing cells, in addition to the endogenous locus^40^. To test fission-line behavior under elevated euchromatic crossover rates, we crossed the lines to the chromosome three arm FTL *420*^30^, which carries markers at base pair positions 260,176 and 5,368,016. To generate a genetic map, the *fission-2* line was crossed to the Ler-0 traffic line *LTL3.4*, with wild type Col-0 used as a control. *LTL3.4* carries fluorescent markers flanking centromere three from base pair positions 10,305,637 to 19,645,919 in the Ler-0 genome and corresponding approximately to chromosome three positions 9,870,598 to 18,429,882 in Col-0 from BLAST. In each FTL line green fluorescence is driven by *NAP:eGFP* and red fluorescence by *NAP:dsRED* transgenes^30,32^.

### Genetic map construction and permutation tests

To generate the genetic map, hemizygous RG/++ F_1_ plants were obtained from the crosses *LTL3.4* (RG/RG)×Col-0 and *LTL3.4* (RG/RG)×*fission-2*. The resulting F_1_ plants were self-fertilized, and 96 F_2_ individuals were collected from a single F_1_ of the Col-0×Ler-0 *LTL3.4* and *fission-2*×Ler-0 *LTL3.4* hybrids. Whole-genome short-read sequencing was performed using 150 bp paired-end reads at a mean coverage of 10× per sample. Samples with evidence of trisomy were excluded, representing 13 individuals from the fission cross and one from the control cross. Reads were mapped using bwa-mem2 (version 2.3) to the Col-0 genome, and biallelic SNPs were called with GATK (v.3.8) using the HaplotypeCaller function^66^. The genetic map was constructed using the R package R/qtl2^67^. SNPs differentiating the Col-0 and Ler-0 genomes were identified using Pannagram^68^, yielding 479,638 SNPs. Biallelic SNPs were retained after filtering for ≤20% missing data and allele frequencies between 0.2 and 0.8, and were thinned to one marker per 10 kb interval to account for markers in linkage disequilibrium, before genetic map construction, finally using 9,493 SNPs.

To test for chromosome-specific differences in crossover frequency, one-sided permutation tests were performed on crossover counts obtained from the genetic map. For each chromosome, the difference in mean crossover number per individual between the *fission-2*×Ler-0 and Col-0×Ler-0 F_2_ populations was calculated, and individual labels were randomly reassigned between populations 1,000 times to generate an expected null distribution. The one-sided *P*-value was calculated as the number of permutations with a difference greater than, or equal to, the observed value.

To examine crossover differences between trisomic and euploid F_2_ *fission-2*×Ler-0 individuals we plotted crossover rate (cM/Mb) for each and observed an increase of crossovers in centromere proximal regions on chromosome 3. Significance was tested by comparing the number of crossovers in the 13 trisomic plants falling within 1Mb of the centromere array compared to 1,000 permutations of euploid F_2_ individuals.

### Segregation distortion analysis

Biallelic SNPs from the Col-0×Ler-0 control F_2_ population and the *fission-2*×Ler-0 F_2_ population were intersected to identify SNPs present in both datasets using bcftools^69^. Shared SNPs were pruned using a 10 kb sliding window, retaining the first SNP per window to reduce linkage disequilibrium. SNP coordinates were lifted over from the Col-0 reference genome to Ler-0 using minimap2 yielding 16,419 matched SNP positions, and all downstream analyses were performed using Ler-0 chromosomal coordinates. Segregation distortion was quantified by comparing SNP allele frequencies between the two populations. Raw genotype counts at each locus were converted to Col-0 and Ler-0 allele counts, and the Ler-0 allele frequency was calculated as the Ler-0 allele count divided by the total allele count at that position. SNPs were divided up into non-overlapping 100 kb windows along the Ler-0 chromosomes. Differences in allele frequency between populations were assessed within each window using a two-sided Fisher’s exact test on the Col-0 and Ler-0 allele counts. The change in Ler-0 allele frequency per window (Δf) was calculated as the difference between the *fission-2* and control SNP frequencies taken from the summed allele counts. Multiple testing was accounted for using Bonferroni correction. For the interval-specific analysis of the *LTL3.4* locus, SNPs within the Ler-0 interval chromosome three from 10,305,637 to 19,645,919 bp (totalling 627 SNPs) were extracted and allele counts were compared in the same manner as genome wide SNPs.

## Notes

### Competing Interest Statement

The authors have declared no competing interest.

